# Rate of meristem initiation driven by the MADS–WUS axis contributes to floral survival and inflorescence evolution in grasses

**DOI:** 10.64898/2026.08.31.748292

**Authors:** Yongyu Huang, Shuangshuang Zhao, Guojing Jiang, Christian W. Hertig, Abdullah Shalmani, Di Peng, Kenan Tan, Twan Rutten, Zihao Zhu, Jinhong Kan, Andreas Maurer, Victor Henrique Rabesquine Nogueira, Amanda Souza Camara, Ping Yang, Klaus Pillen, Ricardo F.H. Giehl, Jochen Kumlehn, Ravi Koppolu, Thorsten Schnurbusch

## Abstract

Crop domestication has repeatedly shaped inflorescence architecture to improve floral production, but mechanisms coordinating the rate of floral initiation, maturation and survival remain unclear. Combining morphometry, modelling and molecular genetic analyses, we show that floral production in the indeterminate barley (*Hordeum vulgare* L.) inflorescence follows an “initiate fast—die young” strategy orchestrated by a main MADS-box gene, *SPIKELET INITIATION AND FERTILITY* (*SIF*). SIF accomplishes this duality by coordinately terminating the inflorescence meristem via *WUSCHEL* and activating the floral meristem via *APETALA1* (*Vrn-H1*). Hereby, the ancestral *SIF* “slow” allele promotes a timely commitment to floral maturation, whereas the derived “fast” allele permits more floral initiations. Postdomestication selection of *SIF* alleles thus enables diversified reproductive strategies in barley populations to maintain yield traits in the field. Finally, we show that a lineage-specific *SIF* duplication contributed to meristem fate transition and inflorescence evolution during *Triticeae* cold adaptation. Our results establish developmental rate as a key driver of architectural innovation and reproductive success.

## Introduction

The evolution of grasses is arguably one of the most underrated success stories especially when one recognizes their importance in terms of ecosystem functions, nutrient cycling, CO2-fixation and food security for our global economies and food demands^1–4^. Grasses are ubiquitous in our daily lives, with gramineous cereal crops such as maize (*Zea mays* L.), rice (*Oryza sativa* L.), wheat (*Triticum aestivum* L.) and barley (*Hordeum vulgare* L.) composing a large proportion of our caloric intake^5^. In particular, during the last 12-2 million years, an unprecedented global cooling fueled a peak in grass diversity that profoundly shaped their life-history strategies, inflorescence architecture and reproductive success^6–8^. These innovations enabled grasses to become ubiquitous across nearly all continents and to dominate ecosystems such as savannas, shrublands, prairies, meadows, pastures and croplands. The Neolithic Revolution, ∼10,000 years ago, marked another milestone in grass evolution, as humans began to domesticate and cultivate cereal grasses, establishing a symbiosis-like relationship that continues to underpin today’s agriculture. Yet despite the ecological and agricultural success of grasses, traits essential for cereal crop domestication and evolution remain only partially understood^9^.

The interconnected traits of lifespan, growth rate and branching define an organism’s fundamental life-history strategy, and have therefore been recurring targets of selection during cereal domestication and evolution^10^. Lifespan (e.g., flowering time) sets the temporal time window for survival and reproduction, whereas branching (e.g., inflorescence branching) influences fecundity and reproduction within plant communities. Growth rate, in turn, governs how quickly individual plants reach their maximal size within a given time window, thereby mediating how effectively plants exploit or allocate resources^11,12^. Despite the central importance of these three life-history traits, they have received unequal scientific attention. For example, while flowering time and inflorescence branching are relatively accessible to measure, and hence, have been extensively characterized across species^13–15^; growth rate has historically been more difficult to quantify due to nonlinear developmental dynamics^16–18^. Our understanding of how growth rate is selected relative to branching during cereal domestication therefore remains limited, leaving its genetic and environmental drivers largely unexplored.

In this work, we sought to systematically dissect the trait growth rate in a crop evolutionary context and examine how it is linked to inflorescence branching. We focus on the temperate *Triticeae* crops barley and wheat, which were key components of the Neolithic Agricultural Revolution. Unlike the multi-branched panicles of rice or the complex tassels and ears of maize, the *Triticeae* inflorescence, called spike, features a less ramified architecture built from a chronologically ordered spikelets initiated along a main axis (rachis)^15^, which enables precise prediction of both spikelet initiation rate and developmental potential (Extended Data Fig. 1A)^15,19^. Despite this conserved architecture, *Triticeae* species differ markedly in the balance between meristem indeterminacy and determinacy. For example, barley maintains an indeterminate inflorescence meristem but highly determinate spikelets, whereas wheat exhibits the opposite tendency. These contrasting developmental outcomes likely reflect differences in how developmental potential is allocated between inflorescence and spikelet meristems, thereby shaping spikelet initiation rate, survival and ultimately inflorescence size. Consequently, spikelets initiated later at the distal end of the spike typically receive less resources, often leading to sterility or degeneration that can eliminate up to ∼50% of total grain yield potential^20,21^ (Extended Data Fig. 1B). Keeping the right pace of spikelet initiation, differentiation and maturation within a confined lifecycle is therefore particularly important for floral production in these annual temperate cereals.

## Results

### An “initiate fast—die young” strategy shapes spikelet initiation and survival in spike-type inflorescences

We reasoned that the rate of meristem initiation in barley is associated with the activity of inflorescence meristems (IMs), pools of stem cells at the growing tips of plants^22^. In many plant species, IM activity is intriguingly linked to its size^23–25^, we therefore fist traced the morphodynamics of barley IM and spikelet meristem (SM) using both light microscopy and confocal laser scanning microscopy. We found that nuclei-occupied cells became coordinately aligned along the IM epidermis at the Waddington 2 (W2) stage^26^, coinciding with the onset of SM emergence (Extended Data Fig. 2A). This morphology represents a maintenance of meristem indeterminacy^27^. Subsequently, deformed nuclei began to appear and were gradually degraded before W5, indicating progressive termination of IM after W2. Concurrently, we observed a linear depletion of IM size along with the initiation of new SMs, consistent with a constant plastochron (the time between two successive SMs) characteristic of alternate-distichous phyllotaxis^19^ (Extended Data Fig. 2B). This further suggests that, although the barley IM is indeterminate, the IM at W2-stage might signify the highest developmental potential for the initiation of SMs. Finally, we assessed whether spikelet development aligns with its timing of initiation and found that proximal spikelets grew faster than distal ones (Extended Data Fig. 2C), indicating that earlier-initiated SMs gain preferential access to resources for differentiation and maturation.

To understand how variation in SM initiation rate (e.g., fast-to-slow) shapes spikelet number, size and survival (defined as the ratio of final spikelet number by maximum spikelet meristems initiated) across different growth durations available for SM initiation, we combined computational modelling with extensive phenotypic dissection (Extended Data Table1; materials and methods). We first considered IM size at the W2 stage as the initial *Developmental Potential* (*DP*), which declines over time along with the initiation of new SMs (Extended Data Fig. 2B). The fate of each SM can be encapsulated with a simple sigmoidal curve^16^, in which the growth rate is a function of its age, or timing of initiation (*t*). *t* varies from *t*_0_ (the onset of initiation) to *t_max_* (the termination of initiation) at a constant plastochron (Δ*t*) (Extended Data Fig. 3A). With these settings, we factorially integrated multiple SM initiation rates with multiple growth duration regimes, thereby generating a two-dimensional theoretical inflorescence morphospace, in which spikelets of varying number and size emerge along a main axis (e.g., the rachis) (Extended Data Fig. 3B,C). We further phenotypically dissected *DP*, Δ*t* and *t_max_* across a set of domesticated barley genotypes (six-rowed spring types, referred to as the D6S panel; n = 358)^28^ and assessed their relationships with spikelet initiation, development, survival, and ultimately, fecundity (Fig. 1A–D; Extended Data Fig. 3D).

**Fig. 1.**
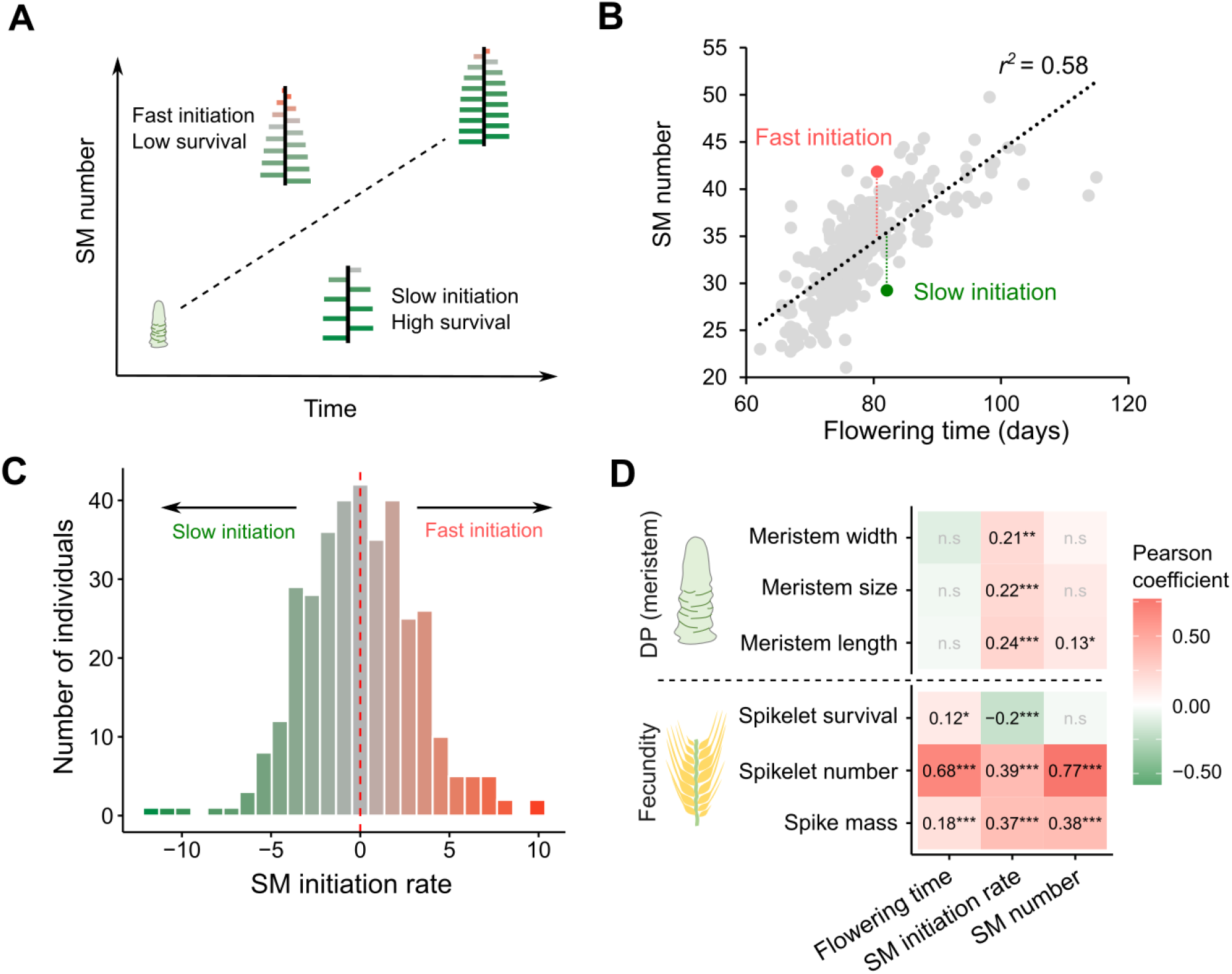
Phenotypic architecture of spikelet meristem initiation rate. (A) Computational modeling for spikelet meristem (SM) initiation, development and survival. Two axes of variation, i.e., growth duration (x-axis) and initiation rate (y-axis), determine lateral organ number. Faster initiation produces smaller distal lateral organs, which are less likely to survive (scaled with red color). (B) Estimation of SM initiation rate in the barley population (D6S, n = 358). Residuals from a linear regression of total growth duration (flowering time) against maximum SM number (recoded as rachis nodes) were used to estimate variation in SM initiation rate. Original data from^28^. (C) A histogram showing the distribution of SM initiation rate in the barley population. (D) Pearson correlation analysis among the three life-history traits, fecundity, and inflorescence meristem size (DP: developmental potential). \*\*\**P* < 0.001; \*\**P* < 0.01; \**P* < 0.05.

Our simulated inflorescences closely recapitulated the architecture of real spikes, including the characteristic proximal–distal decline in spikelet size and maturation. Notably, when a size threshold for survival was applied^15^, the model revealed that faster SM initiation negatively correlates with spikelet size and survival, whereas longer growth durations exert a positive effect (Extended Data Fig. 3C). These simulated architectural outcomes were consistent with empirical phenotypes: larger IM size (*DP*) was associated with faster—and hence more—SM initiations, whereas flowering time (∼*t_max_*) showed a weak, negative relationship with IM size (Fig. 1D). Importantly, although both initiation rate and flowering time contributed positively to maximum spikelet initiation and fecundity, their effects on spikelet survival were opposing, in agreement with model predictions (Fig. 1D).

Taken together, our model and phenotypic data suggest that fast-initiated spikelets tend to be smaller and short-lived; whereas slow-initiated spikelets grow larger and persist longer (Fig. 1A). Moreover, the positive correlation between IM size and SM initiation rate offers mechanistic insights into the developmental basis of SM initiation rate.

### A MADS-box gene controls natural variation in SM initiation rate

To understand the genetic mechanisms of SM initiation rate, we conducted genome-wide association studies (GWAS) using three independent mapping populations (Extended Data Table 2). Population 1 (pop1) comprised the D6S panel (see above); pop2 consisted of 243 backcrossed lines (BC_1_S_3_) selected from the wild barley introgression population HEB-25^29,30^; and pop3 included 130 doubled haploid lines derived from two domesticated barleys (six-rowed, spring-type *vs.* winter-type). Our GWAS consistently revealed a strong association peak for SM initiation rate on the short arm of chromosome 7H across all three populations (Fig. 2A). This locus, designated *SPIKELET INITIATION AND FERTILITY* (*SIF*; named for the Norse goddess Sif, who embodies earth, fertility and fruitful harvests), is syntenic with the top associations for spikelet survival in both pop2 and pop3, but is absent from associations for flowering time or maximum SM number (Extended Data Fig. 4A–F). Moreover, GWAS in pop2 also identified *Photoperiod-H1* (*Ppd-H1*) (Fig. 2A), a gene known to accelerate spikelet initiation but compromise spikelet survival beyond its well-established role as a flowering time promoter^28,31–33^ (Extended Data Fig. 4B,C). Thus, despite being phenotypically interconnected, these three life-history traits—maximum spikelet initiation, flowering time and initiation rate—are controlled by distinct genetic mechanisms.

**Fig. 2.**
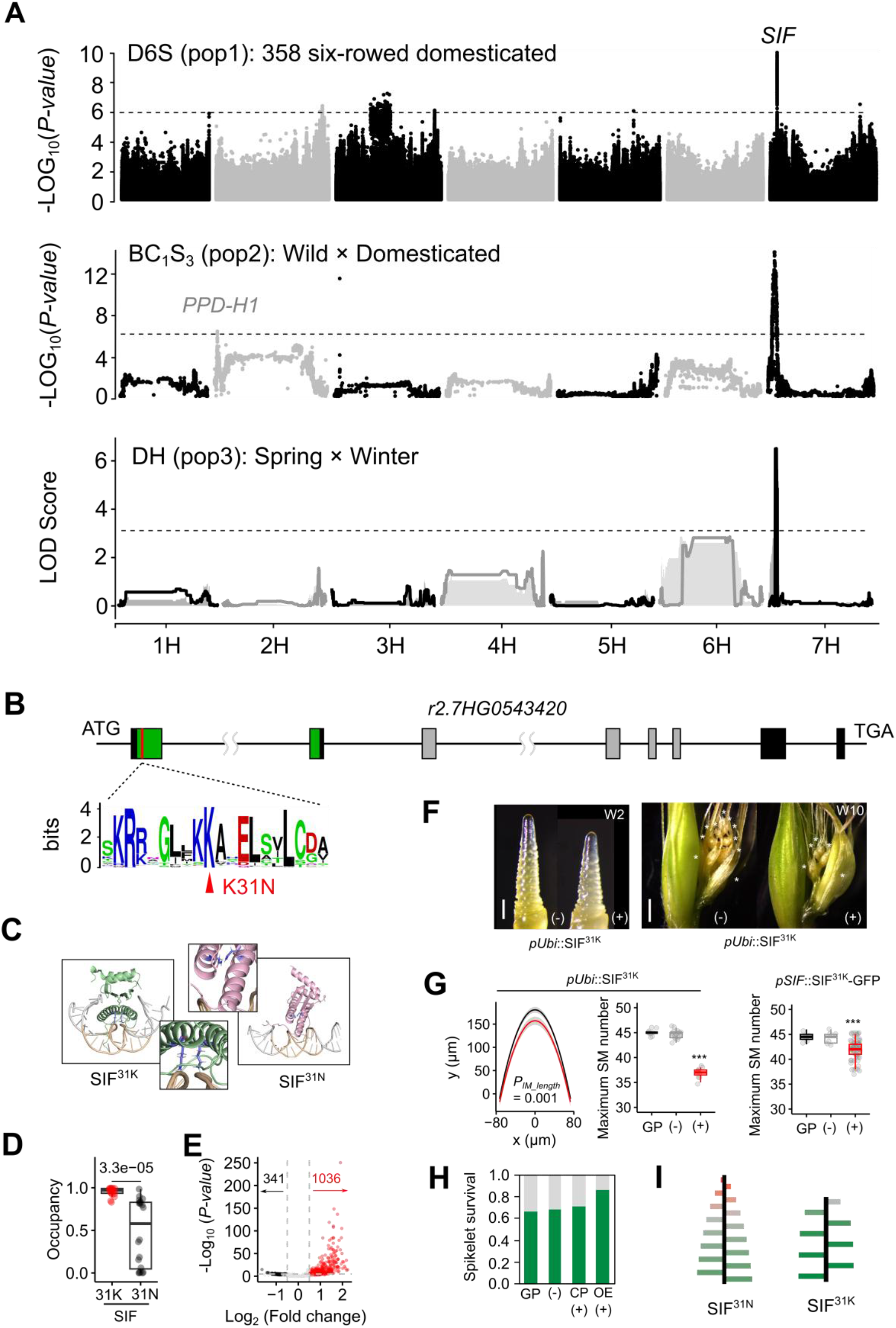
Genetic architecture of SM initiation rate and its functional validation. (A) Genetic architecture of SM initiation rate. Three independent populations converge on a major locus, termed *SPIKELET INITIATION AND FERTILITY* (*SIF*), on the short arm of chromosome 7H. Dashed horizontal lines represent the Bonferroni-corrected significance threshold. (B) Fine-mapping *SIF* to a MADS-box gene. A conserved amino acid substitution (K31N) was highlighted. (C) Functional significance of the K31N substitution in SIF revealed by molecular dynamics simulations. Shown are the final conformations from representative simulations of the SIF^31K^ (green) and SIF^31N^ (pink) protein variants bound to a CArG-box motif from *HvWUS* promoter. DNA sequence is shown in grey (franking)/cyan (core CArG-box) ribbon, and the K31/N31residues is highlighted as blue sticks. Insets show enlarged views of the protein– DNA interaction interface surrounding residue 31. The simulations predict altered DNA-binding interactions between the two SIF variants. (D) The occupancies (fraction of simulation frames) of residues K31 and N31 in interactions with DNA phosphates, for the SIF^31K^ and SIF^31N^ variants. Significant level is determined by a two-sided Student’s *t*-test. (E) A Volcano plot showing a higher enrichment of peaks bound by SIF^31K^ compared with SIF^31N^ in a DAP-seq assay. (F) Functional validation of SIF using genetic transformation. Representative spike images at W2 (left) or W10 (right) stages from transgenic plants are shown. Plants carrying the transgene (+) are compared with non-transgenic segregants (−) from the same transformation event. At W2, pUbi::SIF^31K^ plants exhibit reduced IM size, whereas at W10 they display reduced tip spikelet degeneration. Asterisks indicate residual degenerated floral structures at the spike apex. Scale bars: 200 µm (left panel) and 500 µm (right panel). (G) Statistical comparisons of IM size and maximum SM number from transgenic plants and the background control. Note that *SIF* overexpression (OE; *pUbi*::SIF^31K^) exhibited stronger phenotypic changes in spikelet initiation and survival than plants transformed with the genomic complement construct (CP; *pSIF*::SIF^31K^-GFP). Plants from the T_2_ generation, either with (+) or without (–) the T-DNA insertion, were used for phenotyping. Golden Promise (GP) was used as a control. Significant levels are determined by ANOVA with post-hoc Tukey HSD Test (IM size) or a two-sided Student’s *t*-test (SM number); \*\*\**P* < 0.001. (H) Spikelet survival from transgenic plants and the control. (I) Schematic representation of SIF variation associated with spikelet initiation and survival.

To better understand the developmental basis of *SIF*, we compared meristem states in a pair of *SIF* near-isogenic lines (NIL) derived from pop2 (hereafter the wild barley allele is designated as NIL-SIF^W^, and the cultivar allele as NIL-SIF^C^). NIL-SIF^C^ plants consistently exhibit larger IM size from W2 to W4.5 stage, allowing for faster and more SM initiations compared to NIL-SIF^W^ (Extended Data Fig. 5A–C). Conversely, in NIL-SIF^C^ plants, tip degeneration begins before distal floral identity is specified, leading to reduced spikelet survival (Extended Data Fig. 5D). We found that although the final spikelet number did not differ significantly between NIL-SIF^C^ and NIL-SIF^W^, NIL-SIF^W^ plants tended to exhibit a more uniform grain weight distribution along the spike (Extended Data Fig. 5E), indicating a more balanced resource allocation among spikelets when initiation is slower, consistent with our model and real phenotypic data.

We mapped the *SIF* locus to an ∼150 kb interval containing three annotated genes (Extended Data Fig. 4D–F). Mining DNA sequence variation using multiple reference genomes^34^ identified a single SNP (s7_40905510:C-G) with deleterious impacts (PROVEAN score = -4.681) in *r2.7HG0543420*, a gene encoding a MADS-box transcription factor of the SEPALLATA (SEP) subfamily (named HvMADS5 in^35,36^, hereafter designated SIF). Notably, the rice orthologue *OsMADS5* was shown to control inflorescence branching and SM formation^37^, making *HvMADS5* a good candidate underlying *SIF*. The deleterious SNP, tightly linked to the peak SNP and consistently segregating in the parental lines of our bi-parental mapping populations, results in a lysine-to-asparagine substitution (K31N) at a deeply conserved residue within the DNA-binding MADS-box domain (Fig. 2B). AlphaFold structural prediction followed by molecular dynamics simulations showed that, in the SIF^31K^ variant, K31 maintained stable DNA phosphates interactions for >95% of the simulation time, whereas in SIF^31N^, N31 reached only ∼50% occupancy, suggesting a functional significance of the K31N substitution in DNA binding (Fig. 2C,D). We further validated this using DNA affinity purification sequencing (DAP–seq), which revealed substantially reduced DNA binding capacity of the SIF^31N^ variant compared to the ancestral SIF^31K^ (Fig. 2E; Extended Data Fig. 6A–C; Extended Data Table 3), suggesting that the K31N substitution gives rise to a reduced-function variant of SIF. We validated *r2.7HG0543420* as the candidate gene for *SIF* by complementing the Golden Promise plants (31N) with the 31K coding sequence driven by either a *UBIQUITIN* promoter or the native promoter (Extended Data Fig. 7A). In both assays, transgenic plants exhibited reduced IM size and spikelet initiation, but increased spikelet survival (Fig. 2F–H). Conversely, Cas9-mediated gene knock-out of *r2.7HG0543420* in Golden Promise^36^ did not result in significant phenotypic changes in spikelet initiation or survival (Extended Data Fig. 7B–D), further supporting 31N as a strongly reduced-function protein variant.

Taken together, we conclude that *r2.7HG0543420* is the candidate gene for *SIF*, and that a single amino acid substitution (K31N) attenuates SIF’s function, leading to faster SM initiation and reduced spikelet survival. (Fig. 2I). Notably, the same K31N substitution in another MADS-box gene (*AGAMOUS-LIKE 11*) has been associated with fruit form domestication in oil palm (*Elaeis guineensis*)^38^, suggesting molecular convergence at this substitution.

### SIF-mediated restriction of IM proliferation involves *WUSCHEL*

Our data reveal a dual role for SIF in coordinating IM termination and SM differentiation. In many flowering plants, meristem termination is achieved through the shutdown of *WUSCHEL* (*WUS*), whereas SM identity and subsequent floral organ specification are controlled by the sequential activation and repression of MADS-box genes, including *APETALA1* (*AP1*)^39–41^. Such a developmental coordination stabilizes floral fate and prevents excessive flower production. However, unlike an individual flower, a grass spikelet represents a specialized inflorescence that contains one to several small flowers (florets)^42–44^. In addition, the conservation of *WUS* in organizing stem cells within the grass IMs has been a subject of debate^22,45,46^. We sought to dissect the regulatory logic that enables SIF to couple IM termination with SM activation.

Analysis of tissue-specific RNA-seq data^47^ revealed a spatiotemporal on–off gene expression pattern for *SIF*, *HvWUS* and *VERNALIZATION-H1* (*Vrn-H1*, barley *AP1* homolog), which coincides with the initiation, differentiation and survival of spikelets (Extended Data Fig. 8A,B). For instance, *HvWUS* mRNA levels in both the IM and SM progressively decreases as *SIF* and *Vrn-H1* expression increase. Notably, *SIF* mRNA expression arises in the main spike axis (rachis) after W3 stage when spikelet vascular traces start to emerge^28,48^, suggesting an additional role for *SIF* in vascular development. To investigate the regulatory relationship, we sequenced mRNA from developing spikes in NIL-SIF^C^ and NIL-SIF^W^ plants (Extended Data Fig. 8C,D; Extended Data Table 4). We found that genes related to DNA replication, primordium initiation and IM maintenance (including *HvWUS*) were more highly expressed in NIL-SIF^C^ during the early phase of spikelet initiation. These transcriptomic shifts reflect a delay in SM commitment and prolonged IM activity in NIL-SIF^C^, consistent with its larger IM size and accelerated spikelet initiation, but reduced spikelet maturation and survival. Conversely and consistently, genes associated with photosynthesis, vascular development and floral identity specification (including *Vrn-H1*) were more highly expressed in NIL-SIF^W^ at later developmental stages, resulting in enhanced floral differentiation and survival. Finally, RT–qPCR confirmed the RNA-seq results: *HvWUS* expression was higher in NIL-SIF^C^ at W2, whereas *Vrn-H1* displayed a heterochronic shift, being more highly expressed in NIL-SIF^C^ at W2 but in NIL-SIF^W^ at W4.5 (Extended Data Fig. 9A). These findings are consistent with a role for SIF in promoting SM differentiation.

To test for a direct regulation, we next revisited our DAP-seq data and identified a proximal regulatory region ∼40 bp upstream of the *HvWUS* as a direct binding target of SIF, with notably stronger binding by SIF^31K^ than by SIF^31N^ (Fig. 3A). This region overlaps with an accessible chromatin region identified by ATAC-seq data from leaf tissues^49^ and contains a conserved CArG-box-like motif across temperate grasses of the “core Pooideae” clade^8^, including the Triticeae and Bromeae tribes. The CArG-box is a known binding motif for MADS-box transcription factors and is required for *WUS* expression in different species^50–52^. We first validated the interaction between SIF and the *HvWUS* promoter using an electrophoretic mobility shift assay (Fig. 3B). A dual-luciferase assay in tobacco (*Nicotiana benthamiana*) leaves revealed allele-dependent modulation of *HvWUS* promoter activity, with SIF^31K^ inducing a weaker activation than SIF^31N^ (Fig. 3C). Furthermore, mRNA *in situ* hybridization (ISH) showed that *HvWUS* and *SIF* are expressed in an overlapping region of an organizing center–like domain in the barley IM, albeit *SIF* expression extends further downward within the IM (Fig. 3D). ISH also revealed that both genes are expressed in the SM during its initiation and differentiation (Extended Data Fig. 9B).

**Fig. 3.**
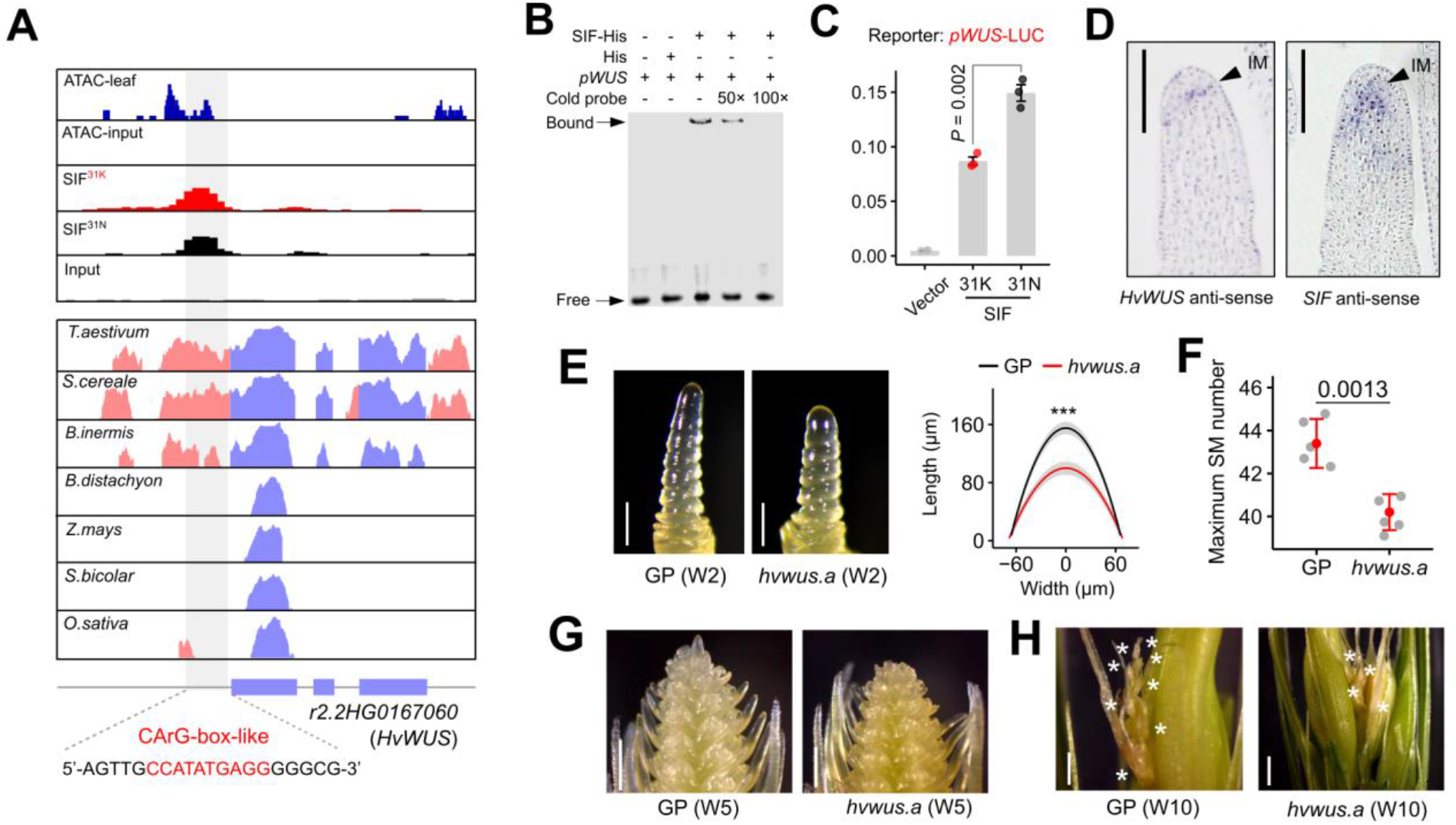
Restriction of IM proliferation by SIF involves *WUSCHEL*. (A) SIF binds to a recently emerged CArG-box-like motif in the *HvWUS* promoter in specific temperate grasses. We identified an overlapping accessible chromatin (top track) in the same region using previously published ATAC-seq data^49^, consistent with this site being a potential upstream gene binding region. The blue and pink colors from the lower box represent conserved genomic sequences from coding (blue) or non-coding regions (pink), respectively. (B) EMSA showing the binding of SIF^31K^ to the conserved CArG-box motif in the *HvWUS* promoter. ‘+’ and ‘−’ indicate the presence and absence of the indicated probe or protein, respectively. (C) Allele-dependent modulation of *HvWUS* promoter by SIF. Heterologous expression of firefly luciferase (LUC) driven by the *HvWUS* promoter was co-transfected with either SIF^31K^ or SIF^31N^ in tobacco leaves. LUC values were normalized to an internal constitutive Renilla luciferase (REN). *HvWUS* promoter co-incubated with an empty vector was used as a control. (D) Tissue-specific expression of *HvWUS* and *SIF* in the inflorescence meristem (IM) during SM initiation using *in situ* hybridization. Both *HvWUS* and *SIF* were expressed in an organizing center–like domain of the barley IM. (E) *HvWUS* positively controls IM size in barley. Note that phenotypes from *HvWUS* weak allele (*hvwus.a*) resemble those in *SIF* WT plants. (F) A comparison of maximum SM number in WT and *hvwus.a*. (G) Representative images showing the distal floral organ differentiation in WT and *hvwus.a* spikes at W5 stage. (H) Representative images showing the distal floral organ differentiation in WT and *hvwus.a* spikes at W10 stage. Asterisks indicate residual degenerated floral structures at the spike apex. Error bars in (C) and (F) represent means ± SD. Significant levels are determined by a two-sided Student’s *t*-test. ***, *P* < 0.001. Scale bars: 200 µm in (D) and (E); 500 µm in (G) and (H).

Finally, we verified the functional relevance of *HvWUS* in IM maintenance and/or SM development using Cas9–mediated mutagenesis. We found that while *hvwus.b* mutant with a frameshift mutation failed to establish an apical meristem and was seedling-lethal (Extended Data Fig. 10), the *hvwus.a* mutant—containing an in-frame substitution of nine consecutive amino acids within the homeodomain of HvWUS— displayed phenotypes resembling those of NIL-SIF^W^ plants, including a smaller IM, reduced SM initiation, and higher survival (Fig. 3E–H). Thus, SIF-mediated restriction of IM proliferation (smaller IM size) involves a timely downregulation of *HvWUS*, which may allow more time for SM differentiation and maturation.

### SIF-mediated SM activation involves a timely activation of *Vrn-H1*

We next interrogated the role of *Vrn-H1* in SM development. Like for *SIF*, ISH revealed that *Vrn-H1* was expressed in developing SMs, albeit with a broader expression extending throughout developing SMs (Extended Data Fig. 9B). This expression pattern prompted us to ask whether elevated *Vrn-H1* expression promotes SM commitment and indeterminacy. In barley, spikelet determinacy is specified by the fate of the spikelet’s axis, the rachilla, which typically produces a single floret followed by meristematic arrest and rapid rachilla elongation (Fig. 4A)^53^. This ultimately culminates in the formation of a sharp tip structure, a feature also observed as the endpoint of indeterminate meristems in other plant species^54,55^. In contrast, the rachilla of rye (*Secale cereale* L.) and wheat remains active and generates additional florets before tip elongation. Thus, an undifferentiated, elongated rachilla reflect reduced SM commitment and premature termination of floral differentiation (Fig. 4A).

**Fig. 4.**
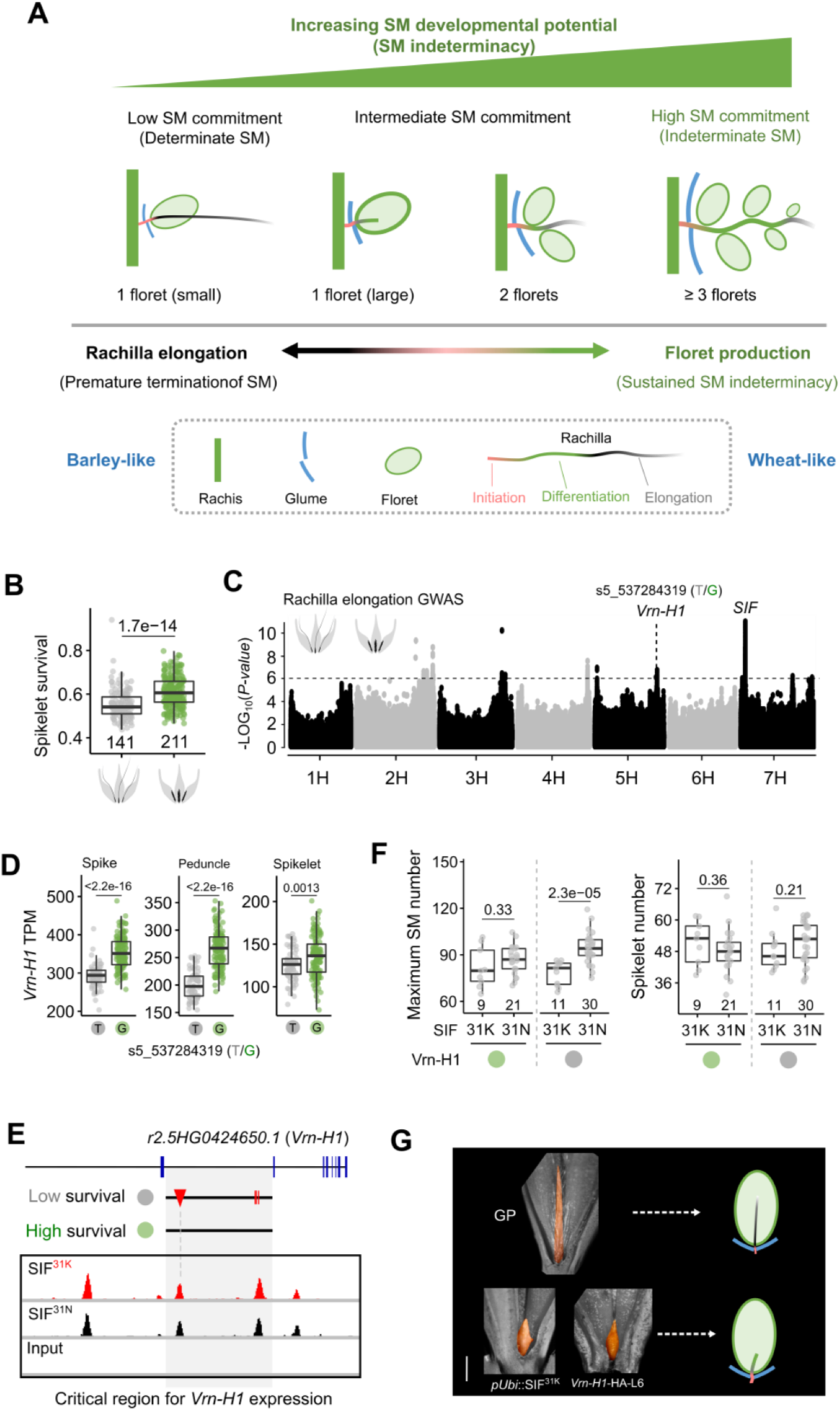
SIF promotes SM differentiation via *Vrn-H1*. (A) Conceptual diagram illustrating a developmental continuum in *Triticeae* spikelet determinacy. Increasing developmental potential promotes sustained SM indeterminacy and continued rachilla differentiation, resulting in the sequential production of additional florets. This also results in enlarged glumes. Conversely, cells undergo rapid elongation if developmental potential is prematurely arrested^19,83^, leading to enhanced rachilla elongation. Colored segments indicate successive developmental states of the rachilla, ranging from initiation (red), differentiation (green), and elongation (grey). The diagram links the transition from barley-like determinate spikelets to wheat-like indeterminate spikelets. (B) Rapid rachilla elongation is associated with lower spikelet survival in the D6S population. (C) Genetic architecture of rachilla elongation. *SIF* remains the main effect, with additional association peaks at the *Vrn-H1* locus tagged by the SNP s5_537284319. (D) The same association peak for rachilla elongation at *Vrn-H1* also explains its expression variation in a two-rowed barley population (see also Extended Data Fig. 11D). Note that the expression association analysis was restricted to three tissues with detectable *SIF* expression (TPM > 1 in ≥50% of genotypes: spike, spikelet, and peduncle). (E) SIF binds *Vrn-H1* based on DAP-seq data. Note that out of the four notable peaks, substantial natural variations, including SNPs (red lines) and large insertions/deletions (red triangle), were identified. See also Extended Data Fig.11F. (F) Epistatic interaction between *SIF* and *Vrn-H1* in pop3. Note that SIF^31K^-mediated repression of SM initiation and promotion of spikelet survival require *Vrn-H1* intron variation. (G) *SIF* and *Vrn-H1* promote rachilla differentiation in transgenic plants. *Vrn-H1* overexpression plants are from^84^. The schematic illustration summarizes the proposed developmental transition from rachilla elongation in GP toward sustained rachilla differentiation and floret production. Scale bar = 500 µm. Significant levels in (B), (D) and (F) are determined by two-sided Student’s *t*-test.

For barley, we consistently found that spikes with elongated rachilla also exhibited lower spikelet survival (Fig. 4B). GWAS on rachilla elongation in pop1 further identified strong association peaks surrounding both *SIF* and *Vrn-H1* (Fig. 4C). We found that the same functional nucleotide polymorphism for *SIF* also explained the genetic association for the elongated rachilla phenotype, with genotypes carrying the SIF^31K^ variant exhibiting shorter rachilla compared to the longer ones in SIF^31N^ (Extended Data Fig. 11A,B). In addition, association peaks (tagged by SNP s5_537284319) at the *Vrn-H1* locus were linked to an intron deletion previously implicated in *Vrn-H1* gene expression^56^, which we found to be in strong linkage disequilibrium (*R²* = 0.94) with the top associations for *Vrn-H1* gene expression (Fig. 4D; Extended Data Fig. 11C–E). Notably, DAP-seq identified four SIF *in vitro* binding sites within *Vrn-H1*, two of which overlap naturally occurring SNPs and insertion/deletion polymorphisms and conferred stronger LUC reporter activation by SIF^31K^ than by SIF^31N^ in transient expression assays (Fig. 4E; Extended Data Fig. 11F,G). Consistently, SIF-mediated inhibition of SM initiation required *Vrn-H1* intron variants in pop3, whereas survived spikelets remained largely unchanged, suggesting epistatic effects of *Vrn-H1* on *SIF* during SM initiation and survival (Fig. 4F). Finally, we found that rachillae in both *SIF^31K^* and *Vrn-H1* overexpression lines were highly differentiated, frequently producing supernumerary floret-like structures (Fig. 4G). Thus, regulation of *Vrn-H1* by SIF promotes SM commitment.

Taken together, we conclude that SIF is a dual-functional transcription factor that coordinately modulates IM activity and SM differentiation, in part via *HvWUS* and *Vrn-H1*. Although direct in vivo binding remains unresolved, our findings providing a developmental framework for the trade-off between rapid spikelet initiation and successful floral maturation and survival.

### Post-domestication selection of *SIF* shaped yield traits across barley row-types

We next examined if SIF-mediated SM initiation rate may be shaped by selection to optimize reproductive outcomes. In barley, the IM produces SM triplets per rachis node. Two-rowed barleys, including all wild progenitors (*H. vulgare* ssp. *spontaneum*), inhibit the two lateral florets during development, yielding only one fertile spikelet per rachis node, whereas six-rowed barleys retain all three florets as fertile (Extended Data Fig. 12A). This morphological transition, i.e., from two-to six-rowed spike row-types, theoretically triples grain yield and is mainly controlled by two transcription factors *VULGARE ROW-TYPE SPIKE 1* (*VRS1*) and *VRS5*^57,58^. Yet, barley plants carrying *vrs1* and *vrs5* mutations rarely reach the expected threefold grain yield increase in the field, likely due to in part a trade-off in SM initiation and survival (Extended Data Fig. 12B)^59,60^.

We first analyzed the frequency of the SIF^K31N^ haplotypes across domesticated and wild barleys^61^, and found that the reduced-function SIF^31N^ variant was completely absent in wild barleys but was progressively enriched in landraces and cultivars (Fig. 5A), suggesting post-domestication selection for SIF^31N^. Notably, the SIF^31K^ variant is present in only ∼3% of domesticated two-rowed barleys, but in up to ∼50% of six-rowed barleys, suggesting that SIF may exert row-type–specific effects on spikelet initiation, survival and/or grain yield. To test this, we compared yield-related traits in two-rowed and six-rowed barleys carrying different SIF variants in field plots (Fig. 5B). We found that, in two-rowed background, despite an increase in spikelet survival in SIF^31K^ genotypes, final spikelet number was reduced due to a substantial reduction in SM number. This indicates that the SIF^31K^ variant exerts negative antagonistic pleiotropy on yield traits in two-rowed barley: while it promotes spikelet survival, it reduces final spikelet number. By contrast, in six-rowed background, SIF^31K^ genotypes showed increases in both spikelet survival and final spikelet number, despite maximum SM number remained largely unchanged. In summary, differential selection of SIF^K31N^ haplotypes fine-tunes reproductive outcomes (spikelet number) across row-types by promoting greater SM initiation in two-rowed barley or enhancing spikelet survival in six-rowed barley.

**Fig. 5.**
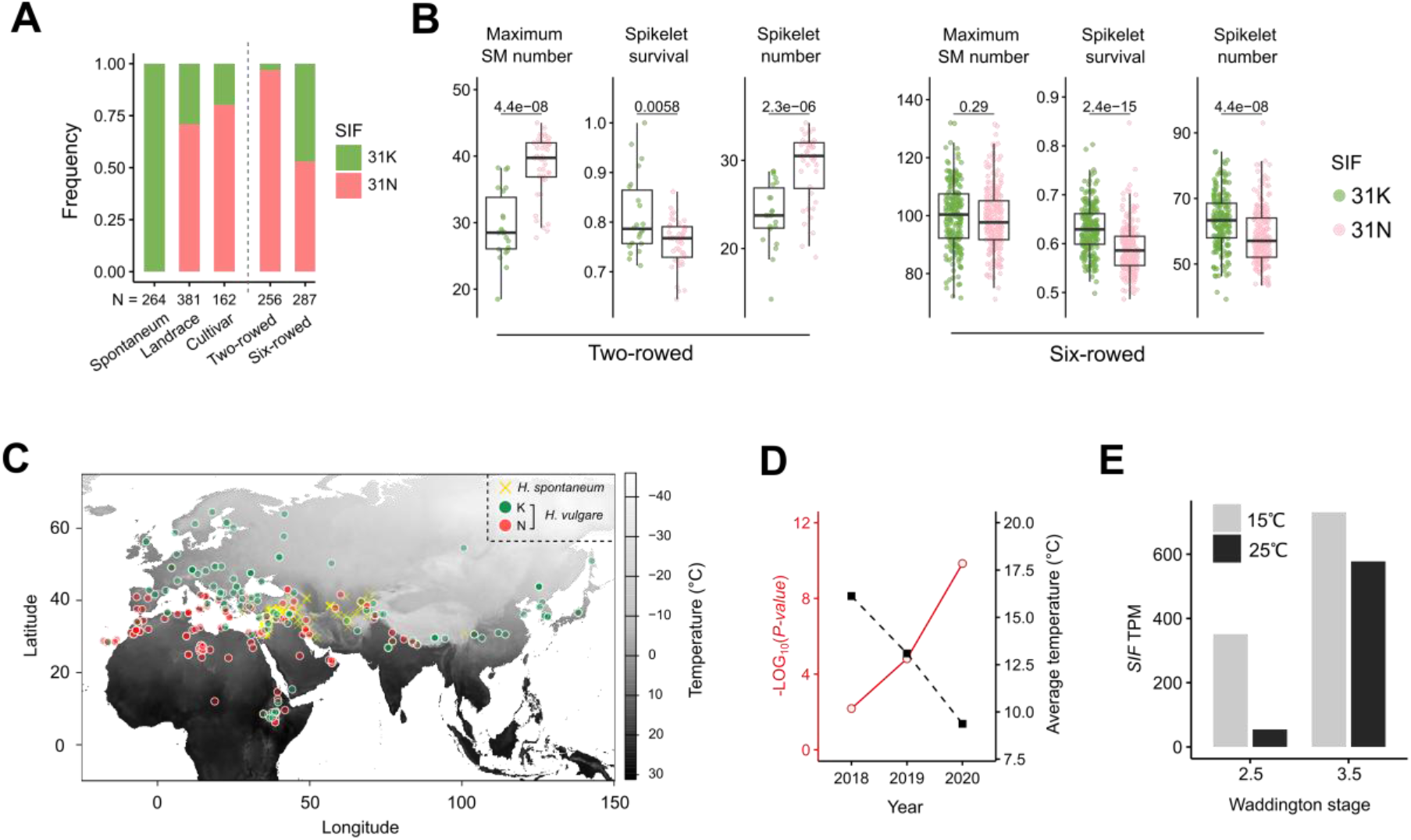
*SIF* selection optimizes reproductive output across row-types during geographic range expansion. (A) SIF^31K^ frequency in wild barleys (s*pontaneum*), early barley domesticates (landrace) and cultivars, as well as different row-types. Source data from^61^. (B) Performance of SIF^K31N^ haplotypes in two-rowed and six-rowed backgrounds in field plots. Two-rowed genotypes were selected from the Genebank of IPK^85^; six-rowed genotypes were from the D6S population. Significant levels are determined by two-sided Student’s *t*-test. (C) Geographical distribution of barley with different SIF variants (264 wild accessions, 287 six-rowed accessions). Mean annual temperature data from WorldClim (https://www.worldclim.org/) were projected onto the map. (D) GWAS of SM initiation rate across three years (2018–2020) in common garden experiments at IPK Gatersleben. SIF’s genetic effect on SM initiation rate was negatively correlated with average temperature. Source phenotypic data from^20^. See also Extended Data Fig. 14. (E) High ambient temperature (25 °C) represses *SIF* expression in developing spikes under Golden Promise (SIF^31N^) background. Source data from^36^.

### *SIF* selection correlates with environmental adaptation across a temperature gradient

To gain insights into how SIF haplotypes arise along environmental gradients, we examined the selection footprints of SIF haplotypes across domesticated and wild barleys^34^. Analysis of genetic diversity (π) first revealed an ∼60-kb region surrounding SIF with substantially lower diversity in SIF^31N^ genotypes than in SIF^31K^ genotypes or wild barleys (Extended Data Fig. 13A). The localized reduction in diversity suggests a selective pressure targeting the SIF^31N^ haplotype, presumably because it conferred a yield advantage by promoting faster and more SM initiations. A median-joining haplotype network and a visualization of genomic structure further revealed independent selections of SIF functional haplotypes (K31N) during barley crop evolution (Extended Data Fig. 13B,C). SIF^31N^ haplotype featured a single origin, with the most closely related wild counterparts found in Israel; whereas SIF^31K^ arose at least twice during the separation of Eastern and Western domesticated barleys. Mapping the demographic distribution of SIF haplotypes using a geo-referenced barley panel^61^ revealed a clear geographic cline for SIF haplotype frequency (Fig. 5C). Fast-initiation haplotypes carrying the reduced-function SIF^31N^ variant predominated in southern latitudes with high annual temperatures, whereas SIF^31K^ frequency increased toward northern and eastern regions with colder climates. The positive correlations between SIF “fast–slow” haplotypes and “high–low” ambient temperature are in broad agreement with a pan-latitudinal gradient in the global fast–slow continuum of plant life-history strategies^62^. We thus suggest temperature as a key environmental driver shaping *SIF* haplotype diversity to optimize reproductive success during barley range extension. Consistent with this idea, we observed a negative correlation between *SIF*’s genetic effect on SM initiation rate with local temperature in a three-year field trial (2018-2020)^20^, and that *SIF* expression in developing spikes was down-regulated at higher temperatures (Fig. 5D,E; Extended Data Fig. 14).

In summary, our results suggest multiple origins of SIF haplotypes in domesticated barleys, and highlight temperature as a key environmental driver of their selection, consistent with previous population genetics studies^63,64^.

### *SIF* duplication drives an evolutionary shift in meristem determinacy in *Triticeae*

Given that the diversification of *Triticeae*—including barley, wheat and rye—over the past 12–5 million years occurred during a period of global cooling^8^ (Fig. 6A), we reasoned that natural selection at *SIF* may have contributed to lineage-specific floral innovation by facilitating adaptation to colder environments. A key developmental hallmark of this adaptive evolution is the transition in meristem determinacy within the spike inflorescence. For instance, each SM produces a single floret in barley, 2–3 in rye and up to 12 in wheat; by contrast, the barley IM exhibits greater indeterminacy, rye is intermediate, and wheat’s IM terminates early into a SM (Fig. 6B). This evolutionary transition appears to be accompanied by an enlargement of glumes, which presumably canalizes meristem identity towards SM fate, thereby promoting the production of multiple florets within a spikelet^65,66^. We observed a similar enlargement of the glumes into lemma-like structures in *SIF* overexpression barley plants, and consequently, spikelets occasionally (∼10%) produced additional florets (Fig. 6C). Moreover, in certain recombinants carrying the SIF^31K^ variant from our bi-parental populations, distal spikelets survived after heading, with some even terminating their IM into a spikelet (Fig. 6C).

**Fig. 6.**
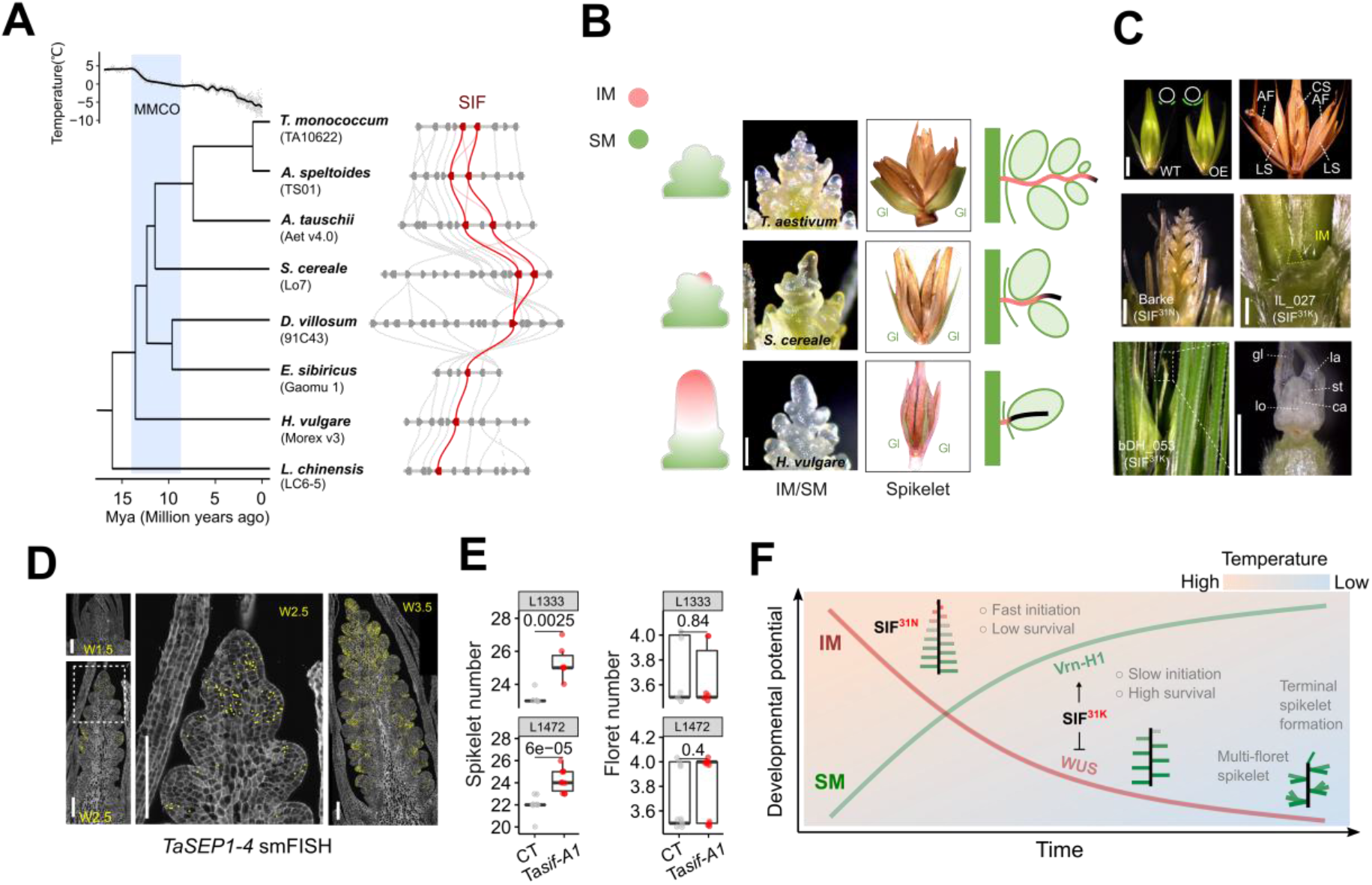
SIF duplication drives Triticeae spike evolution. (A) Syntenic block of the *SIF* region across the genomes of nine species, representing approximately 16 million years of Triticeae evolution since the Middle Miocene. The *SIF* duplication occurred after the speciation of the *Dasypyrum* genus (*D. villosum*) and coincided with the Mid-Miocene Climatic Optimum (MMCO), a period of global cooling. Phylogeny three and divergent time of the species were extracted from^8^. Syntenic block was built under the database (https://wheat.cau.edu.cn/TGT/)^86^. (B) A gradual shift in meristem determinacy during the evolution of Triticeae spikes. Representative species from barley, rye, and wheat were selected (middle). Schematic illustrations depicting the gradual loss of IM indeterminacy and the concurrent gain of SM indeterminacy are shown on both sides. (C) Loss of IM indeterminacy and gain of SM indeterminacy driven by SIF^31K^ functionality. Enlargement of the glume and the emergence of multi-floret spikelet phenotypes were observed in SIF^31K^ overexpression lines. Certain recombinants carrying the SIF^31K^ variant in populations pop2 and pop3 exhibit 100% spikelet survival at heading or develop terminal spikelet phenotypes. gl, glume; lo, lodicule; la, lemma; st, stamen; ca, carpel. (D) Expression of wheat *SIF* orthologs (*TaSEP1-4*) across two spike developmental stages. Gene expression was marked with yellow spots. Source data were based on previously published single-molecule fluorescence *in situ* hybridization (smFISH) data^87^. (E) *TaSIF-A1* mutant spike phenotype. Sibling plants segregating from two different *TaSIF-A1* TILLING lines (Cadenza background) were used to assess spike morphology. CT denotes non-mutant sibling control plants, whereas *Tasif*-A1 represents plants carrying the corresponding *TaSIF*-A1 mutation. Significant levels are determined by two-sided Student’s *t*-test. (F) Conceptual model illustrating the gradual shift in developmental potential between the IM and SM during *Triticeae* spike evolution and temperature adaptation. The SIF^31N^ allele promotes IM maintenance and rapid spikelet initiation, resulting in higher spikelet number but reduced spikelet survival. In contrast, SIF^31K^ represses *WUS* and activates *Vrn-H1*, promoting developmental potential allocation to SMs, slower spikelet initiation, and increased spikelet survival. Progressive shifts in meristem commitment are proposed to underlie the transition from indeterminate barley-like spikes to determinate wheat-like spikes with multi-floret spikelets. The blue gradient indicates the proposed effect of temperature, from high (left-top) to low (right-bottom), on developmental potential allocation medited by the MADS (SIF –VRN-H1)–WUS regulatory module. Scale bars: 200 µm in (B) and (D); 500 µm in (C).

We then examined whether the SIF-mediated meristem determinacy pathway may have contributed to morphological diversification of Triticeae spikes using phylogenetic analyses. As reported previously^67^, the SEP-subfamily MADS-box genes, including the *SIF* ortholog, are duplicated in wheat compared to rice, and we found that this duplication occurred after the divergence of *Hordeum* within the *Triticeae* (Fig. 6A; Extended Data Fig. 15A,B). Spatial gene expression data^68^ showed overlapping wheat *SIF* (*TaSIF*) expression in the IM, SM and rachis, as compared to barley (Fig. 6D), indicating a conserved function in spikelet initiation and differentiation. Consistently, loss of its closest ortholog, *TaSIF-A1*, in hexaploid wheat increased spikelet number per spike without altering flowering time, while floret number per spikelet remained largely unchanged (Fig. 6E; Extended Data Fig. 15C), suggesting enhanced IM activity and accelerated spikelet initiation. Previous reports showed that *AP1* was duplicated more than once before the emergence of the *Pooideae* and likely contributed to the formation of the grass spikelet^8,44^. Based on these findings, we propose that the duplication of *AP1-*/*Vrn1*-homologs^8^, followed by *cis*-regulatory gain leading to down-regulation of the central meristem homeostasis gene *WUS*, positioned SMs at the two flanks of the IM (alternate-distichous phyllotaxis) in Triticeae inflorescences. A subsequent *SIF* duplication in the wheat lineage further repressed the IM and promoted SM development; these stepwise events may have driven the evolution of Triticeae spikes across different temperature ranges and thereby contributed to their today’s diverse inflorescence patterning (Fig. 6F).

## Discussion

Here, by combining large-scale phenotyping with modeling and molecular genetics, we established a conceptual framework to understand how growth rate (rate of SM initiation) acts in concert with flowering time to shape inflorescence size and adaptive evolution. Mechanistically, we identified a meristem determinacy pathway involving two MADS-box genes and the key meristem homeostasis gene *HvWUS*, which together regulate meristem size, rate of SM initiation and spikelet survival.

In plants, larger meristems are thought to initiate lateral organs faster by providing additional physical space for organ initiation, potentially facilitating the synchronized differentiation and maturation of organs^19^. Consequently, targeted manipulation of developmental genes such as those in the CLAVATA–WUS pathway, which control meristem size, has been shown to influence yield-related traits in many species^50,53,69,70^. A further reduction in meristem size may shift determinacy by replacing the apical meristem with the most recently formed lateral organ (terminal flower formation), as demonstrated in barley and other species across a broader evolutionary span^71,72^. In barley and other *Triticeae* species with smaller IMs, it remains unclear how changes in meristem size could improve yield performance due to their constrained inflorescence morphospace (e.g., lateral floral organs can only emerge sequentially on opposite sides of the IM; alternate-distichous phyllotaxis). Indeed, our findings, together with a previous study^73^, suggest that both large and small IMs can be beneficial for yield traits in this temperate cereal grass, and that subtle shifts in meristem size encoded by a single SNP in *SIF* can enhance either floral initiation or floral survival to maintain yield traits. SIF encodes a SEP MADS-box gene; members of which were initially identified as floral organ identity regulators^74^, and subsequently found to control broader developmental processes such as inflorescence branching^36,37,75,76^, grain size^77^ and fruit ripening^78^, suggesting their broad ecological and agricultural implications.

Our study highlights a complex interaction between flowering time and meristem initiation rate that collectively shapes overall plant size. For example, while both *Ppd-H1* and *Vrn-H1* were initially identified as main flowering time promoters^33,79^, both exert opposite effects on the rate of SM initiation, with *Ppd-H1* acting as an accelerator and *Vrn-H1* serving as a stabilizer of floral meristem fate. Nevertheless, the prevalence of mixed *SIF* allelic combinations with flowering time genes during crop evolution illustrates how diversifying combinations of life-history traits (rate and lifespan) via breaking co-adapted wild gene complexes can reschedule the pace of reproduction during range expansion. Beyond controlling organ initiation, both *SIF* and *Vrn-H1* are homeotic genes known to operate in a complex regulatory hierarchy during the speciation of different floral organ identities^80^. Indeed, whereas both *SIF* and *Vrn-H1* are progressively upregulated in the IM during SM initiation, their biphasic expression dynamics in SMs (Extended Data Fig. 8B) may suggest a delayed negative-feedback model in which SIF initially enhances Vrn-H1 activity to specify SM and rachilla fate, but subsequently attenuates it to promote later SM differentiation. Although interactions between SEP-and AP1/VRN1-class proteins are well established in meristem-fate specification^40,81,82^, our findings add additional layers of regulatory complexity for these MADS-box genes. Future studies linking SIF to different functional variants of *Vrn-H1* (e.g., promoter and intron) will provide further insights into how this complex regulatory relationship shapes the evolution of reproductive strategies and will be directly relevant for optimizing crop performance.

## ACKNOWLEDGEMENT

We thank J. Dubcovsky for sharing the *TaSIF* smFISH data and the critical comments, B. Trevaskis for sharing the *Vrn-H1* transgenic materials, D. Zhang for sharing the *SIF*-edited transgenic materials in Golden Promise background, N. Stein and M. Mascher for pre-publication access to barley resequencing data, A. Himmelbach for NGS service on the D6S population, C. Hartmann, Z. Guo and E. Chen for sharing Triticeae spikes for photographing, I. Otto and J. Schippers for assisting protoplast isolation and luciferase measurements, H. Ma and L. Zhang for suggestions on the *Triticeae* phylogenomic analysis, T. Venkatasubbu, R. Kamal and N. Shanmugaraj for assisting barley phenotyping, A. Fiebig for data submission, S. Sommerfeld for barley transformation, C. Trautewig, K. Wolf, A. Püschel, A. Ziplys and I. Otto for technical support, E. Geyer and P. Schreiber for plant care, all members from the PBP and FPS groups for fruitful discussions.

## Authors contributions

Conceptualization: Y.H. and T.S.; Methodology: Y.H.; Investigation: Y.H., S.Z., J.G., C.W.H., D.P., K.N., T.R., Z.Z., V.-H.-R.N., A.-S.C., A.M., K.P., J.K., P.Y., M.M., J.D., J.C. R.K. and T.S.; Visualization: Y.H.; Funding acquisition: Y.H. and T.S.; Supervision: Y.H. and T.S.; Writing – original draft: Y.H.; review and editing: all authors.

## Data and materials availability

RNA-seq and DAP-seq data are available from the European Nucleotide Archive (https://www.ebi.ac.uk/ena/browser/home) under the accession numbers PRJEB97709 and PRJEB97708, respectively. Source code for simulating proximal-to-distal floral organ size can be found on Zenodo (https://zenodo.org/records/17377029). All other data are available in the main text or the Extended Data materials. All materials are available upon request from the corresponding author.

## Materials and Methods

### Plant materials, growth conditions and phenotyping

#### Plant materials

The three mapping populations used in this work are from previous studies^28,30^. 358 six-rowed domesticated spring barleys (pop1: D6S) were selected from the IPK Genebank^85^; they all carry a sensitive allele of *Photoperiod-H1* (*PPD-H1*) based on SNP22 located in the CCT domain^33^. The 25 wild barley founder parents, together with the 247 introgression lines (pop2: BC_1_S_3_), are from a previous research^29^. The 130 doubled haploid (DH) lines were produced by crossing BCC149 (winter barley) and BCC719 (spring barley). Two-rowed barleys with SIF^31K^ allele were selected from the IPK Genebank^85^. Specifically, among the 20,458 accessions in the whole collection, 38 were identified as spring-type, two-rowed domesticated barleys carrying the sensitive *PPD-H1* allele [selection marker: s2_29202906 (v1), r² = 0.97 with SNP22 at *PPD-H1*], which we confirmed with a caps (cleaved amplified polymorphic sequence) marker according to the SIF functional nucleotide polymorphism (G93C – K31N). Near-isogenic lines (NILs) differing at the *SIF* locus were generated using a heterogeneous inbred family (HIF) analysis approach^88^ by marker-assisted selection of homozygous lines from segregating BC_1_S_6_ population (HEB_19_027 descendants, see also Extended Data Fig. 4E. Wheat *TaSIF-A1* mutants (WCAD1333 and WCAD1472, heterozygous) in *cv.* Cadenza background^89^ were ordered through the UK Germplasm Resource Unit (GRU) (https://www.seedstor.ac.uk/shopping-cart-tilling.php). Homozygous sibling lines were then selected for phenotyping following a similar HIF approach. Barley early maturity mutants were ordered from the NordGen seed bank. Golden Promise (two-rowed, spring type) was used as the recipient for genetic transformation. Barley *VRN-H1* overexpression lines (VRN-HA-L6) are from a previous study^84^. All plant materials used in this study are summarized in Extended Data Table 2.

#### Growth conditions

For estimating meristem initiation rate, plants were cultivated under either glasshouse conditions (photoperiod: 16 h/8 h, light/dark; temperature: 20 °C / 16 °C, light/dark. Pop1-3, in 9-cm^2^ pots) or the field station (pop1, 2018-2022, ∼300 plants/m^2^) at the IPK according to^20,28,30^. For transcriptomic study and meristem morphometry, we grow plants under a controlled phyto-chamber conditions (photoperiod: 12 h / 12 h, light / dark; temperature: 12 °C / 8 °C, light / dark), which was also used in our previous floral transcriptomic study^47^. For measuring grain weight distribution from SIF NILs spikes, plants were grown under field conditions at the IPK Lage station, with twelve plants per row (row width: 1.2 m) and a row spacing of 10 cm.

#### Phenotyping

Flowering time (FT; referred to here as *t_max_* in the model), maximum spikelet primordium number (MYP; recorded as maximum rachis node number), final spikelet number (SN, recorded as final rachis node number) and spikelet survival were collected previously^20,28,29^. Other phenotypes, including spikelet initiation rate, meristem size (referred to here as DP, or developmental potential, in the model), rachilla elongation and spike biomass were deduced or collected from this study. Details of all phenotypic data are summarized in Extended Data Table 3.

All inflorescence-related phenotypic data were collected from the main culm. Maximum spikelet primordium number was determined according to^26,90^. Final spikelet number was counted between heading and grain filling stage and spike biomass was measured after harvesting. Four to five randomly selected plants were used for every trait to phenotype the 358 barleys (pop1), the 130 DH (pop2). Barley accessions were grown in a completely randomized design to reduce residual variances. We obtained the individual phenotypic variance components by using a linear mixed-effect model. For spikelet-tracking experiments, barley mutants and corresponding wild-type were dissected every 2-3 days under a light-sheet microscopy. For meristem morphometry, images taken by AxioVision (SE64 Rel. 4.9.1) were directly measured in the same software. Spikelet survival was deduced by dividing SN by MYP^28^. Spikelet meristem initiation rate was estimated by extracting residuals from a linear regression of MYP with flowering time. Due to the variability in developmental timing, W2 stage–specific meristem size variation was successfully captured in 241 of the 358 accessions from the D6S population. All phenotypic data analyses were performed in R (R-3.6.1).

### Modelling spikelet initiation, size and survival

We modeled the size of each spikelet based on its timing of initiation from the inflorescence meristem (IM). The model was based on our detailed observations of the morphodynamics of early barley spike development, along with the reasonable assumption that finite resources are allocated among offsprings (developing spikelets) to maximize fitness (i.e., trade-offs between offspring number and size)^91^. That is: (i) the indeterminate barley IMs implicates infinite spikelet initiation, yet spikes from different barley genotypes consistently reach their maximum spikelet number at a certain stage called “awn primordium” stage (W4.5), suggesting feedback signals from the developing spikelets that restrict further spikelet initiations; (ii) size of the apical meristem declines linearly along with the initiation of new SMs, indicating a depletion of resource from the apical meristem for spikelet initiation; (iii) earlier-initiated spikelets exhibit higher growth rate compared to the distal, younger ones, indicating competition for resources among spikelets during development.

The SM number *N*(*t*) at a specific time (*t*) can be modeled as:

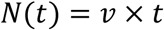

Where *v* is spikelet initiation rate (spikelets per unit time), which is inversely related to a plastochron.

The growth rate of different spikelets (*b*) can be modeled as:

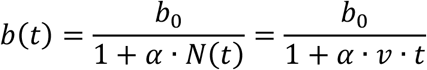

Where *b*(*t*) is the effective growth rate of spikelet primordia at time *t*; *b*_0_ is the baseline growth rate when few spikelet primordia exist (e.g., reduced competition); *α* is the strength of inhibition from existing spikelets (e.g., with competition). This is because, when more spikelets are initiated over time, they compete for limited resources for growth, thereby reducing the growth rate of the spikelets. In this scenario, faster initiation (higher *v*) would give rise to more SM number *N*(*t*), and therefore slower growth rate of the initiated spikelets.

For a given spikelet (or any case of organogenesis) under finite resources, its ontogenetic trajectories (*S*), can be described with a simple sigmoidal curve^16^: first increase gradually (e.g., W1.5 – W2, primordia formation), more rapidly in the middle phase (e.g., W2 – W4.5, floral organ differentiation), and slowly at the end, levelling off at a maximum size after a certain developmental stage (e.g., W4.5 – afterwards). We incorporated our observation that each spikelet’s growth rate is correlated with its initiation timing, and modeled spikelet size dynamics (*S*) as a time-dependent growth function:

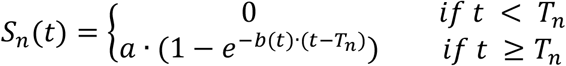

Where *S_n_*(*t*) is size of the *n* − *t*ℎ spikelet primordium at time *t*; *T_n_* is the initiation time of the *n* − *t*ℎ spikelet primordium, which is equivalent to 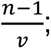 *a* is the maximum spikelet size at awn primordium stage; *b*(*t*) is the time-varying growth rate at time *t*. Further desperations of the modelling parameters were given in Extended Data Table 1.

We ran the simulation by factorially integrating multiple SM initiation rates *v* ∈ [*v*_0_, *v_max_*] with multiple growth durations *t* ∈ [*t*_0_, *t_max_*]. In the model, growth duration is a theoretical parameter describing the duration of spikelet initiation and serves as a proxy for flowering time; accordingly, longer growth durations correspond to later-flowering genotypes. This generates a two-dimensional morphospace of spike architectures with varying spikelet number and size. We assume that a spikelet can survive only if its final size exceeds a specified threshold, such that spikelets failing to reach this minimum size fail to survive. Simulation was performed in R (R-3.6.1).

### Marker-trait association studies

Marker-trait association studies were performed with the same settings according to^28,30^. Briefly, GWAS in the D6S (pop1) was done by considering a set of 22,405,297 bi-allelic SNPs. We used a linear mixed model that incorporated pairwise genetic similarities (kinship matrix) as the random effect and additional population structure informed by a principal component analysis (PC1–5) as the fixed effect (Q+K model). We ran the GWAS using the software GEMMA^92^, with a less stringent significant cutoff at *P* < 1 × 10^−5^. Linkage disequilibrium (LD) was analyzed using PLINK^93^ to identify SNPs in the genomic region surrounding a specific leading SNP using the --r2 command. GWAS in the wild barley introgression lines (pop2, BC_1_S_3_) was done using a simple linear regression model by considering the family information as a covariate under GAPIT3^94^. We considered a significant cutoff as *P* < 1 × 10^−3^, which was similar to a previous research^95^. QTL analysis in pop3 was done with QTL ICiMapping v4.2^96^. Inclusive composite interval mapping was used (ICIM-ADD method) to detect QTL. Population type was set as F1DH (F1-derived doubled haploids). Permutation tests (1,000 times) were used to determine the significance threshold at *P* < 0.05. Association mapping for *Vrn-H1* expression variations in 209 two-rowed barleys was performed using GEMMA with the same Q+K model by considering the first three PCs as the fixed effect. Gene TPMs (Transcripts Per Million) across three tissues (spikelets, peduncle, spikes) were extracted from^97^. These tissues exhibit notable *SIF* expressions, although SIF FNP was almost fixed in this population. All genome-wide association plots were generated using CMplot (https://github.com/YinLiLin/CMplot).

### Barley transformation

For Cas9-mediated mutagenesis of *HvWUS*, two target motifs were selected with CRISPR-P 2.0 (http://crispr.hzau.edu.cn/CRISPR2/) using the *HvWUS* genomic sequence from Golden Promise. A cognate gRNA was considered suitable if it (i) showed specificity to the target gene as verified by BLASTn, and (ii) lacked substantial secondary structure formation within its target-specific 20-bp 5’ end, as predicted using the RNAfold webserver (http://rna.tbi.univie.ac.at/cgi-bin/RNAWebSuite/RNAfold.cgi). The *cas9*/gRNA construct was generated using the CasCADE modular vector system^28^. For the *SIF* overexpression assay, the 678-bp SIF coding sequence from a wild barley parent (K allele) was PCR-amplified and cloned into the UbiFull-AB-M vector with the maize *Polyubiquitin 1* (*Ubi1*) promoter. For the SIF genetic complementation assay, the same *SIF* coding sequence (without a stop codon) was cloned into the pGFP_Amp vector, with the GFP coding sequence fused to the 3′end of SIF (vector 1). Then, the *SIF* promoter (3,427 bp) from the same wild barley parent was PCR amplified and inserted at the 5′end of *SIF* in vector 1, resulting the *pSIF*::SIF^31K^-GFP expression cascade (vector 2). Finally, all expression units (*cas9*, gRNA and *SIF* overexpression) were mobilized into binary vector p6-d35S-TE9 (DNA Cloning Service) via Sfi I restriction digest and ligation. Resultant vectors were used for *Agrobacterium*-mediated T-DNA transfer into immature embryos of barley cv. Golden Promise^98^. Following regeneration under hygromycin selection, transgenic plantlets were screened for the presence/absence of T-DNA insertion. Target region-specific PCR amplicons of were then subjected to Sanger sequencing for detection of locally induced mutations. Primers used for vector construction are summarized in Extended Data Table 6.

### Structural modelling for SIF DNA binding

Protein structures were predicted with a local installation of AlphaFold2^99^. For the molecular dynamics setup, we used PyMOL^100^ to position the protein ∼10 Å away from the DNA along the z-axis, based on the protein–DNA orientation in PDB ID 3KOV as a reference, and whose DNA molecule sequence we mutated to the CArG-box we found upstream *HvWUS* (Fig. 3A). Surface charge distributions analysis was performed using the APBS plugin from PyMOL^101,102^. Structural figures were created with PyMOL software.

The molecular dynamics systems were prepared using the CHARMM-GUI Solution Builder^103–105^. Protein–DNA complexes were parameterized with the CHARMM36m force field^106^. Each system was solvated with explicit TIP3P water molecules, and K^+^/Cl^−^ ions were added to reach a physiological salt concentration of 0.15 M KCl, ensuring charge neutrality. The solutes were placed in rectangular simulation boxes with a 12.0 Å buffer from the edges. Periodic boundary conditions were applied.

To ensure robust sampling, we carried out 20 independent MD simulations for each of the two systems. All simulations were performed in NAMD 3.0^107^ and followed a three-step protocol: minimization, equilibration, and production. During minimization, each system was relaxed for 10,000 steps using the conjugate gradient method. This was followed by equilibration in the canonical (NVT) ensemble for 1 ns at 303.15 K, maintained with a Langevin thermostat (damping coefficient = 1 ps^-1^).

The production phase consisted of 200 ns unrestrained MD in the isothermal– isobaric (NPT) ensemble at 303.15 K and 1 atm. Temperature was controlled with a Langevin thermostat, while pressure was regulated using a Langevin piston barostat (damping time = 1 ps^-1^, oscillation period = 50 fs). A 2-fs integration time step was applied, and the SHAKE algorithm was used to constrain all bonds involving hydrogen atoms. Simulation trajectories were subsequently analyzed and visualized in PyMOL. Using MDAnalysis package^108,109^, we calculated occupancy as the fraction of simulation frames in which residue 31 was contacting any DNA phosphate atoms by less than 4 Å.

### DNA affinity purification sequencing (DAP-seq)

DAP-seq library construction and sequencing were done at SeqHealth (China) Company Limited. Briefly, SIF (HvMADS5) coding sequences from a wild barley (31K, WT allele) and Barke (31N, reduced-function allele) were polymerase chain reaction (PCR) amplified and fused with a SP6 promoter at the 5’ end and a 3x FLAG tag at the 3’ end. *In vitro* protein production was done using a TnT^®^ SP6 High-Yield Wheat Germ Protein Expression System (Promega L3260) according to the manufacturer’s instructions. 6 μg input DNA template was used in a 50μl TnT reaction. After 2 h incubation at room temperature, the 50 μl TnT reaction was mixed with 20 μl anti-FLAG magnetic beads (Merck Millipore). Following 1 h incubation at room temperature, the anti-FLAG magnetic beads were immobilized, and washed three times. The genomic DNA of Morex developing spikes (∼Waddington stage 4.5) was prepared using a Plant Genomic DNA Kit (Tiangen Biotech., Beijing, China). 1 μg of genomic DNA was sheared to an average length of 300-500 bp via ultrasonication for 16 min at 20% amplitude with 2 s on and 8 s off on ice. The sheared DNA fragments were ligated with Illumina adaptor sequences using a VAHTS Universal DNA Library Prep Kit for Illumina V3 (Catalog NO. ND607, Vazyme, Nanjing, China), and named “input”. Anti-FLAG beads with bound SIF^31K^ and SIF^31N^ were each incubated with 50 ng input library. Then the DNA of affinity purification was extracted by phenol-chloroform method and named “DAP”. The DNA fragments were PCR amplified, purified, quantified and finally sequenced on Novaseq 6000 sequencer (Illumina) with PE150 model. Sequencing reads were first filtered by Trimmomatic (version 0.36)^110^ and mapped to the reference genome of Morex version2^111^ using bowtie2 (version 2.2.6)^112^. Peak calling and annotation were done using the MACS2 software (Version 2.1.1)^113^ and the bedtools (Version 2.25.0)^114^, respectively. Motifs enrichment analysis was done using Homer (version 4.10) with the “findMotifsGenome.pl” function^115^. All bound peaks and the adjacent genes identified in this work were summarized in Extended Data Table 3.

### Electrophoretic mobility shift assay (EMSA)

The SIF coding sequence from a wild barley (31K) was cloned into pCold-TF to generate an N-terminal His-tag fusion and expressed in E. coli BL21(DE3). His-TF-SIF was purified using Ni–NTA affinity chromatography, eluted with imidazole, and dialyzed into binding buffer (20 mM HEPES pH 7.5, 50 mM KCl, 5% glycerol, 1 mM DTT). Cyanine 5.5 (Cy5.5)-labeled oligonucleotide probes containing the CArG-box motif in the HvWUS promoter were synthesized, annealed, and incubated with purified His-SIF in 20 μL binding reactions using the Odyssey® EMSA Kit (LI-COR Biosciences, cat# 829-07310), containing 50 ng/μL poly(dI–dC) and 5% glycerol at room temperature for 30 min. DNA–protein complexes were resolved on 6% native polyacrylamide gels in 0.5× TBE at 4°C, and analyzed with the LI-COR Odyssey Imaging System. Specificity was assessed via competition with excess unlabeled cold probe. Probes and primers for EMSA were given in Extended Data Table 6.

### In planta protein–DNA interaction assay using dual luciferase reporter system

We PCR amplified the *HvWUS* promoter region (∼1500 bp) and three SIF-bound cis-regulatory fragments (peal1-peak3, 900-1000 bp) from *Vrn-H1*, and cloned into the pGreenII 0800-LUC vector^116^ to generate the reporter constructs. Full-length coding regions (678 bp) of *SIF* from wild barley (31K allele) or Barke (31N allele) were cloned into the pGreenII 62-SK vector equipped with a 35S promoter (effectors). Nicotiana benthamiana plants were grown under 16 h light / 8 h dark at 22–24 °C and leaves from 4–6-week-old plants were used for transient expression. The above vectors were introduced into Agrobacterium tumefaciens GV3101, cultured overnight, and resuspended in infiltration buffer (10 mM MES pH 5.6, 10 mM MgCl₂, 150 µM acetosyringone) to an OD₆₀₀ of 0.4–0.6. Bacterial suspensions were mixed and pressure-infiltrated into the abaxial side of leaves using a needleless syringe. Leaf tissue was harvested 48–72 h post-infiltration, frozen, and extracted in passive lysis buffer before clarification by brief centrifugation. Luciferase and renillia luciferase (REN, for normalization) activities were detected with a Dual-Luciferase Reporter Assay System (Promega, E1910) under the GloMax Discover plate reader system from Promega. Primers used for the dual-LUC assays were summarized in Extended Data Table 6.

### RNA-sequencing

#### Sample collection and RNA extraction

Developing spikes from NIL-SIF^C^ and NIL-SIF^W^ plants grown under phyto-chamber conditions (see above) were hand-dissected under a light microscope and pooled into 2.0 mL Eppendorf tubes pre-cooled in liquid nitrogen. Samples were always collected at the same time slot (11:00 to 14:00) during a day, and 5 – 20 samples were pooled to make one replicate, depending on the tissue amounts. Total RNA was extracted with TRIzol (Invitrogen) and precipitated with 2-propanol. Genomic DNA residues were removed with a TURBO DNA-free™ Kit (Invitrogen, AM1907).

#### Sequencing and DEG analysis

Library construction and sequencing were done at Novogene (UK) Company Limited. Briefly, Total mRNA was enriched with oligo(dT) beads and randomly fragmented with fragmentation buffer. A stranded specific library was then prepared for each sample and sequenced with Novaseq 6000 PE150 platform, yielding about 50 million high-quality reads per sample. To quantify gene transcripts, RNA-seq reads were first trimmed for TruSeq3-PE adaptors with Trimmomatic v.0.39^110^ using a maximum of two seed mismatches, a palindrome clip threshold of 30, a simple clip threshold of 10, and reads shorter than 36 bp were removed. We used Morex genome annotation V2 as a reference to estimate read abundance with Kallisto software^117^ by using strand-specific mode with the first read reversed (--rf-stranded) parameter. Gene-level abundance (transcripts per million, TPM) and counts were summarized with tximport (v3.14)^118^, and were given in Extended Data Table 4. Only genes with read counts ≥10 in at least 3 samples were retained for DEG analysis under DESeq2^119^. Gene with |log2 FC| ≥ 1 or |log2 FC| ≥ 0.5 (for W2 stage) in expression, and a Benjamini-Hochberg FDR-adjusted *P-value* < 0.05 was considered as a DEG, resulting in 2,539 DEGs caused by *SIF* variation, as summarized in Extended Data Table 5.

#### Clustering and functional enrichment analyses

Average TPM values from the identified DEGs were normalized and subsequently clustered using the K-medoids method with the PAM algorithm, as implemented in the R package cluster (v2.1.3)^120^. We used Euclidean distance as the dissimilarity metric and determined the number of partitions to be clustered based on Gap-Statistics, resulting in six functional clusters (C1 – C6). For GO term enrichment analysis, we used the corresponding *Arabidopsis* homologs, defined as the genes with the best BLASTp hits (e-value < 1e−5). GO term enrichment was done with Metascape (http://metascape.org)^121^ using default parameters and summarized in Extended Data Table 5. Representative genes from each functional enrichment category (C1 to C6) were highlighted in Extended Data Fig. 8D.

### mRNA in situ hybridization

For probe preparation, gene specific (*Vrn-H1*, *HvWUS* and *SIF*) fragments (∼500 bp) were PCR amplified using the total cDNA of BW spikes and cloned into the pGEM-T cloning vector. Vectors confirmed with sanger sequencing were used as templates for the preparation of sense (negative control) and antisense probes. A fusion primer set containing a 20-bp T7 promoter sequence (5’-TAATACGACTCACTATAGGG-3’) before the forward primers of sense probes or reversed primer of antisense probes were used to generate templates for in vitro reverse transcription with T7 RNA polymerase. Spikes were first hand-dissected and fixed overnight with FAA (50% ethanol, 5% acetic acid and 3.7% formaldehyde) at 4℃. Samples were dehydrated with series of ethanol (50, 70, 85, 95 and 100%) and then embedded into Paraplast Plus (Kendall, Mansfield, MA). Sections (8 µm) were prepared using a microtome and mounted onto Superfrost plus slides. Tissue pre-treatment, hybridization, washing and coloration were performed according to^122^. Primers for amplifying the probe sequences were summarized in Extended Data Table 6.

### RT-qPCR

Total RNA extraction and the removal of genomic DNA were done as mentioned in the RNA-seq section. First-strand cDNA was synthesized from total RNA using the SuperScript III Reverse Transcriptase Kit (Invitrogen, 18080-051) according to the manufacturer’s instructions. Quantitative real-time PCR (RT–qPCR) was performed using SYBR Green Master Mix (Thermo Fisher Scientific, A46112) on an PTC Tempo 384 Thermal Cycler (Bio-Rad). Relative transcript abundance was calculated using the comparative Ct (2^−ΔΔCt) method and normalized against the reference gene *HvActin*. Primers used for RT–qPCR are listed in Extended Data Table 6.

### Microscope imaging

For scanning electron microscopy (SEM), spike samples were fixed in 50 mm cacodylate buffer (pH 7.2) containing 2% glutaraldehyde and 2% formaldehyde at 4 °C. Samples were washed with distilled water and dehydrated in an ascending ethanol series and point-dried in a Bal-Tec critical point dryer (https://www.leica-microsystems.com). Dried specimens were gold-coated in an Edwards S150B sputter coater (http://www.edwardsvacuum.com) and examined in a Zeiss Gemini30 scanning electron microscope (https://www.zeiss.de) at 10 kV acceleration voltage. Fixation, resin embedding, sectioning, histological staining and imaging of developing apexes from *hvwus.b* and WT were performed as described previously^123^.

### Phylogenetic and haplotype analysis of SIF and related MADS-box genes

For the chosen eight Triticeae species, their phylogenomic positions were taken from^8^. The global temperature curve over the last 16 million years were modified from^124^. Collinearity and duplication of SIF in Triticeae was conduct at https://wheat.cau.edu.cn/TGT/ ^86^. We reconstructed a phylogenetic tree for the LOFSEP clade of SEPELATA MADS-box genes, with an emphasis on Triticeae species. Three grass-specific members (MADS1, MADS5 and MADS34) from rice and barley were queried against proteomes from 10 species downloaded from Phytozome v12.1 (https://phytozome-next.jgi.doe.gov/)^125^, which resulted in 71 genes. Phylogenetic trees were built with RAxML^126^. We carried out rapid bootstrapping and best-scoring ML tree searching in the same run (-f a) with an extended majority rule (- # autoMRE). The resulted tree was visualized with iTOL (https://itol.embl.de/).

To investigate sequence variations at the *SIF* locus, SNPs within a 1 Mb interval surrounding the SIF gene a previously reported 100 wild and 200 domesticated barleys^127^. A neighbor-joining clustering based on the distance matrix was done with PHYLIP 3.68 (https://evolution.genetics.washington.edu/phylip). The output tree was visualized with ggtree^128^ in R (R-3.6.1).

---

**Extended Data Fig. 1.**
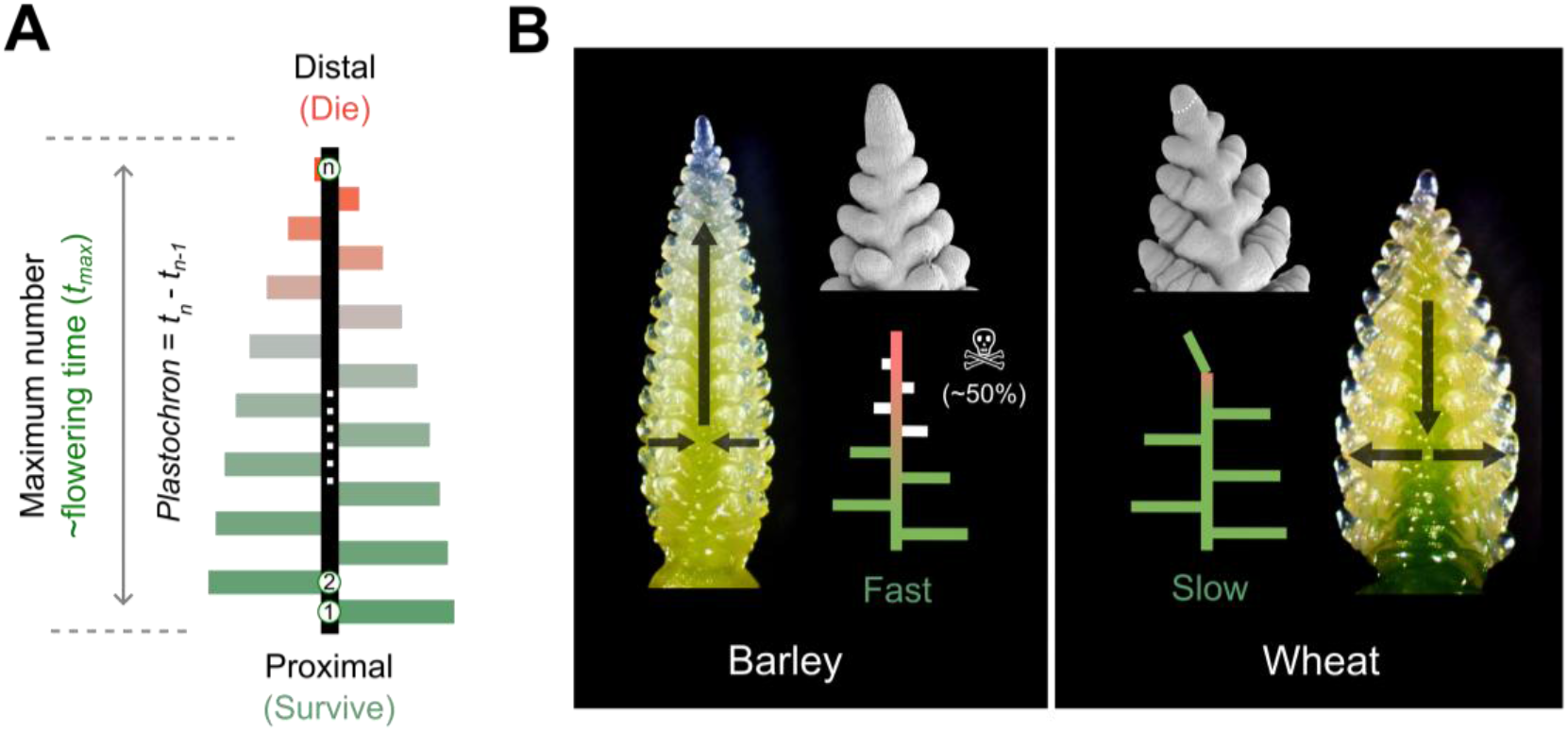
The alternate-distichous organization of spikelets and the proximal–distal decline in spikelet maturation, size and survival. (A) Schematic illustration of the alternate-distichous phyllotaxis in Triticeae spikes. Each module, or spikelet, is illustrated with a horizontal line that aligns along a main axis (rachis, in black) from proximal to distal. In each iteration, the size (scaled with green to red) of the newly initiated spikelet declines relative to the previous one at a predictable mathematical pattern (see also Extended Data Fig. 3). The total duration required to initiate the maximum number of spikelet meristems is linearly correlated with flowering time, and the time interval between two adjacent spikelets is referred to as a plastochron. This architectural design principle is intrinsically connected to the gradient in spikelet maturation, size and survival. (B) Developing barley and wheat spikes at comparable developmental stages (∼W4.5). Scanning electron microscopy (SEM) images reveal a larger IM in barley than in wheat. Consequently, barley is proposed to allocate more developmental potential toward lateral organ initiation and thus exhibit faster SM initiation, but reduced distal SM differentiation, ultimately leading pre-anthesis tip degeneration (marked in skull) that can eliminate up to 50% of initiated SMs^20,28^. Wheat exhibits the opposite pattern: a smaller IM that ultimately terminates as an SM (white dashed outline in the SEM, accompanied by increased developmental potential allocated to individual SMs, which become relatively indeterminate and produce multiple florets. Grey arrows indicate the proposed direction of developmental potential allocation and meristem commitment between the IM and SMs.

**Extended Data Fig. 2.**
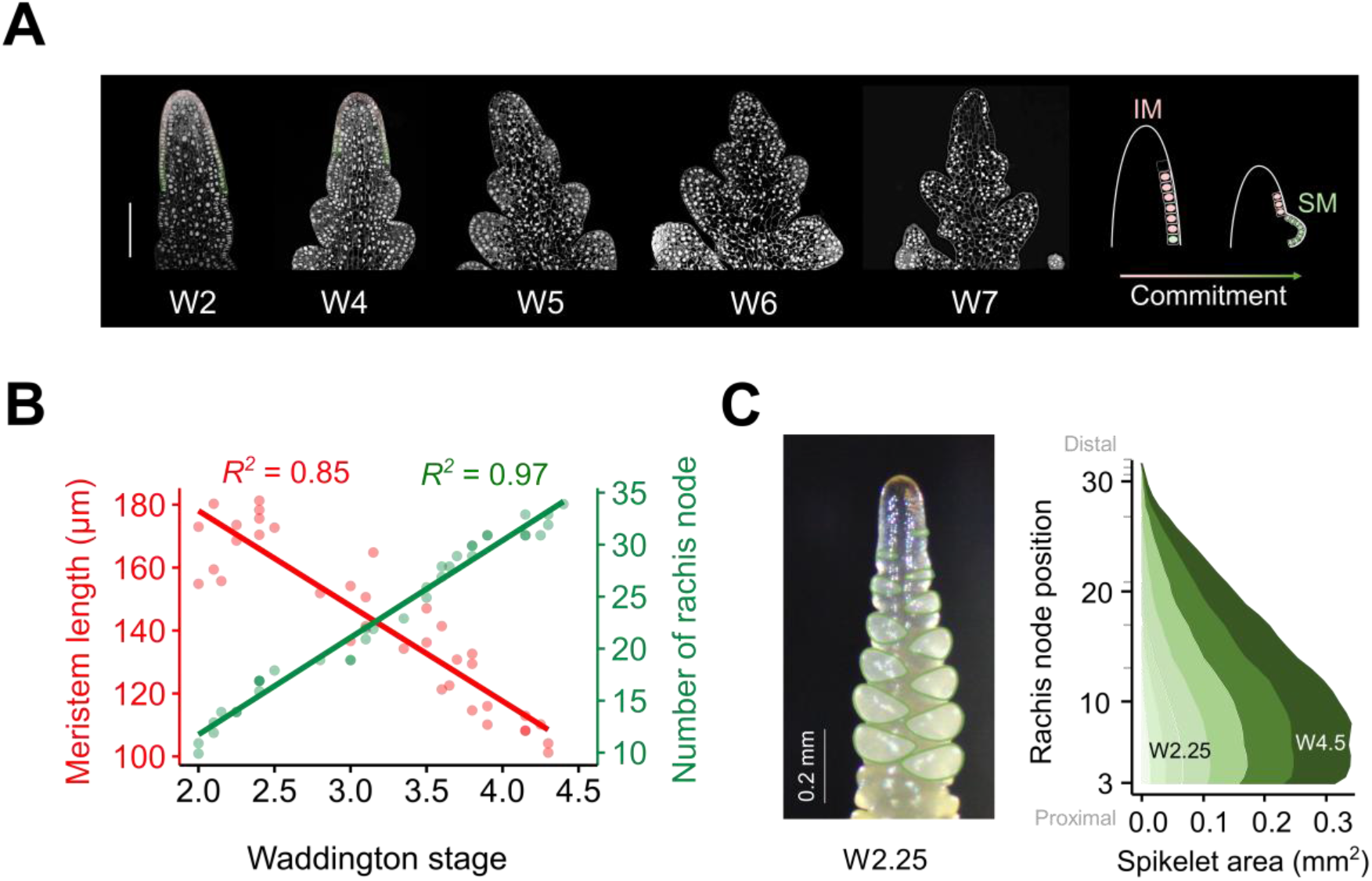
Morphodynamics of IM and SM in barley. (A) Morphodynamics of the barley inflorescence meristem (IM) over time revealed by confocal laser scanning microscopy of longitudinal sections from Waddington stage 2 (W2) until W7. Barley developing spikes were embedded, sectioned and stained with 4’,6-diamidino-2-phenylindole (DAPI). The transition from aligned (W2) to irregular (W4 and afterwards) epidermal cell organization marks a structural manifestation of meristem termination before visible pre-anthesis tip degeneration at ∼W7. A schematic illustration of IM size reduction during SM initiation was shown on the right. Scale bar: 100 µm. (B) IM size decrease along with the initiation of new SMs from W2 until W4.5 in a WT genotype Bowman. Linear fits of developmental stages vs. IM size (red) or SM number (green) were included. Note that the SM number is equivalent to three times the number of rachis nodes. (C) Spikelet growth along the spike axis assessed by changes in spikelet area in a WT Bowman. Left, representative spike at W2.25 with individual spikelets outlined in green. Right, changes in spikelet area along the proximodistal axis between W2.25 and W4.5. Each shaded profile corresponds to a Waddington developmental stage, with lighter to darker shades representing progressively later stages. Grey ticks on the y-axis indicate the latest rachis node positions present at the corresponding Waddington stage, reflecting the progressive increase in spikelet number during spike development. Proximal spikelets are not only initiated earlier but also grow faster compared with the distal ones (see the model).

**Extended Data Fig. 3.**
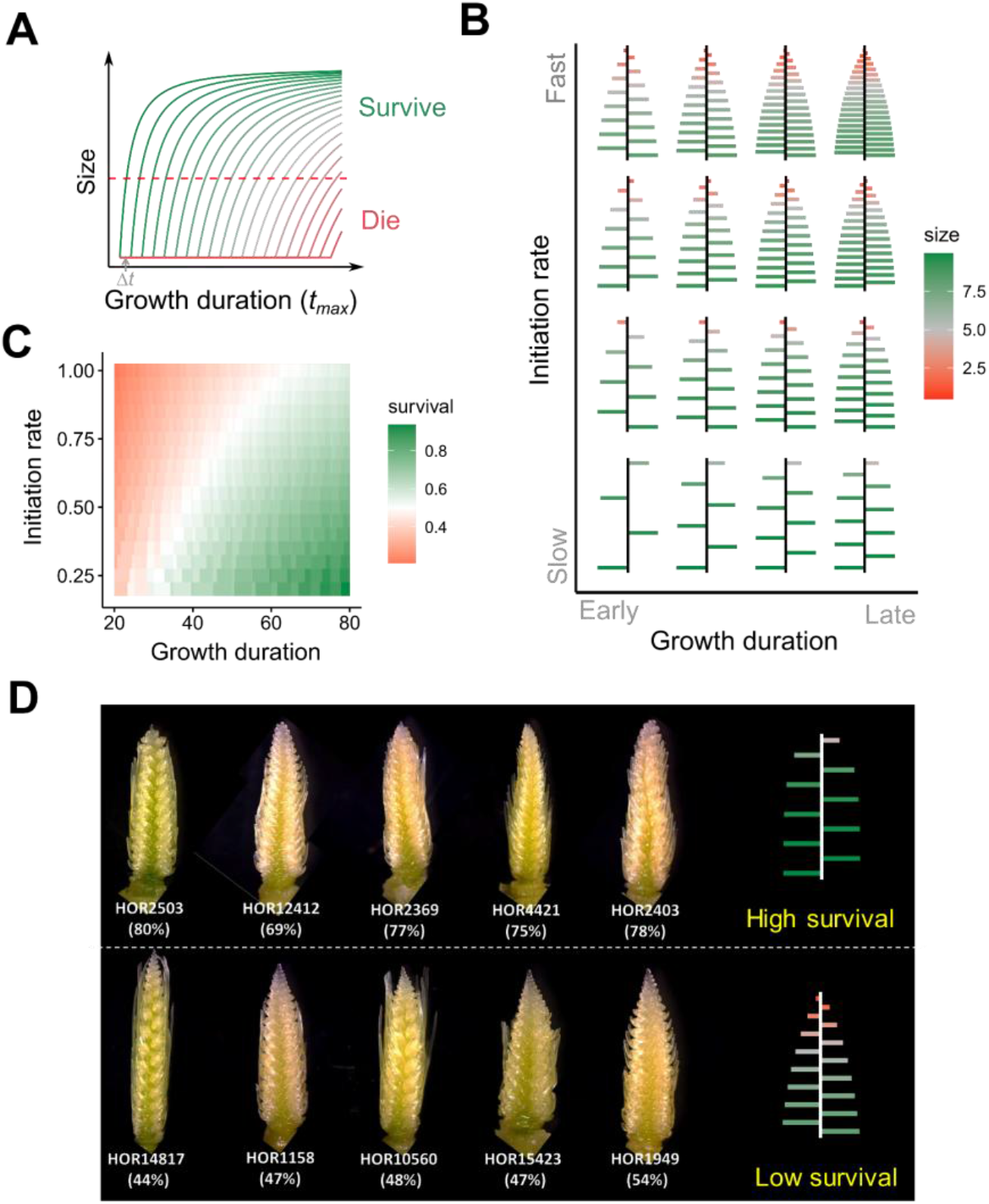
Simulated and real inflorescences illustrating natural variation in SM initiation, development and survival. (A) Asymptotic curves illustrating lateral organ size dynamics in relation to their timing of initiation. Within a given growth duration, lateral organs initiated at a certain plastochron (Δ*t*) are shown in green, with subsequent development following logistic growth at distinct rates (slope of the curve). Red dashed lines denote a predefined size threshold required for survival. (B) Simulated inflorescences with different growth durations and lateral organ (spikelet) initiation rates. Lateral spikelet size was plotted against its timing of initiation. Faster initiation produces more lateral spikelets but results in a steeper proximal-to-distal developmental gradient (scaled with the red-green gradient). (C) Heatmap illustrating how spikelet survival varies with different initiation rates and growth durations based on simulations. Note that growth duration shows a positive correlation, whereas initiation rate shows a negative correlation (scaled with the red-green gradient). (D) Representative spike images from genotypes with low and high spikelet survival. In low-survival genotypes, the IM collapses earlier, and the distal floral organs are smaller and less differentiated compared to those in high-survival genotypes (see also Fig. 1D). HOR numbers represent different barley genotypes. Percentages indicate spikelet survival.

**Extended Data Fig. 4.**
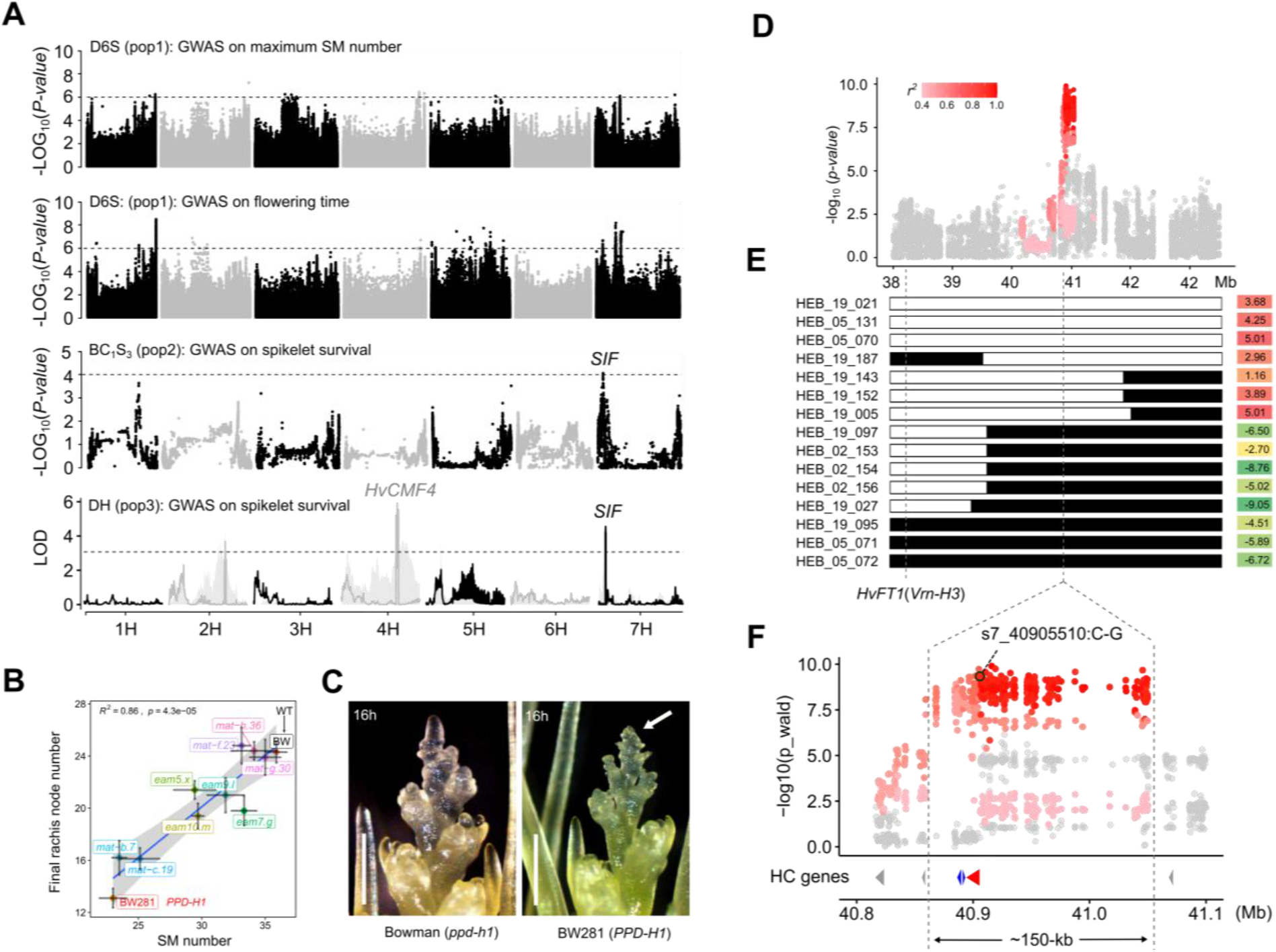
Association mapping of additional life-history traits and the fine-mapping of *SIF*. (A) GWAS for SM number, flowering time and spikelet survival. Note that GWAS did not reveal the same association peak for maximum SM number and flowering time as *SIF* at chr7H, indicating that these three life-history traits have largely independent genetic bases. In contrast, the same association peaks for *SIF* were detected for spikelet survival in pop2 and pop3, supporting a functional link between spikelet initiation rate and survival. Moreover, GWAS for spikelet survival in pop3 also identified a previously characterized CCT MOTIF FAMILY gene (*HvCMF4*)^28^. GWAS for flowering time (pop1) and spikelet survival (pop3) were reproduced from our previous data^28,30^. Dashed horizontal lines represent the Bonferroni-corrected significance threshold. (B) *Ppd-H1* accelerates spikelet initiation but decreases spikelet survival under long-day conditions (16h). Regression of final rachis node number with maximum SM number among ten different early maturity mutants. Note that while all early maturity (*eam* or *mat*) mutants exhibit less maximum SM number compared to the WT Bowman (BW, *ppd-h1* allele), only BW281 (*Ppd-H1* allele) and *eam7.g* close or outside the lower edge of the confidence interval (95%), indicating reduced spikelet survival in both lines. Error bars represent means ± SD. (C) Representative images showing the early tip degeneration in BW281 compared to control (Bowman) under long-day conditions. Images are taken from a same experiment previously^28^. Scale bars: 500 µm. (D) Local Manhattan plot at the *SIF* locus. SNPs were colored according to their linkage relationship (*r^2^*) with the lead SNP. (E) Recombinant mapping narrowed *SIF* down to an ∼150 kb genomic interval using the HEB wild barley introgression lines. The genomic location of a flowering-time gene, *VERNALIZATION-H3* (*Vrn-H3* or *HvFT1*), was indicated, located ∼2 Mb downstream of the candidate gene (*HvMADS5*). Recombinants were sourced from a previously developed wild barley nested association mapping population^29^, with their spikelet initiation rates (fast vs. slow) indicated on the right (see also Extended Data Table 2). (F) Zoom-in to the mapping interval. High-confidence genes in the mapping interval are colored with blue and red (*HvMADS5, SIF* candidate) according to Morex V2 annotation. Position of the SNP (s7_40905510:C-G) that leads to an amino acid substitution (K31N) in HvMADS5 (red arrowhead) was highlighted.

**Extended Data Fig. 5.**
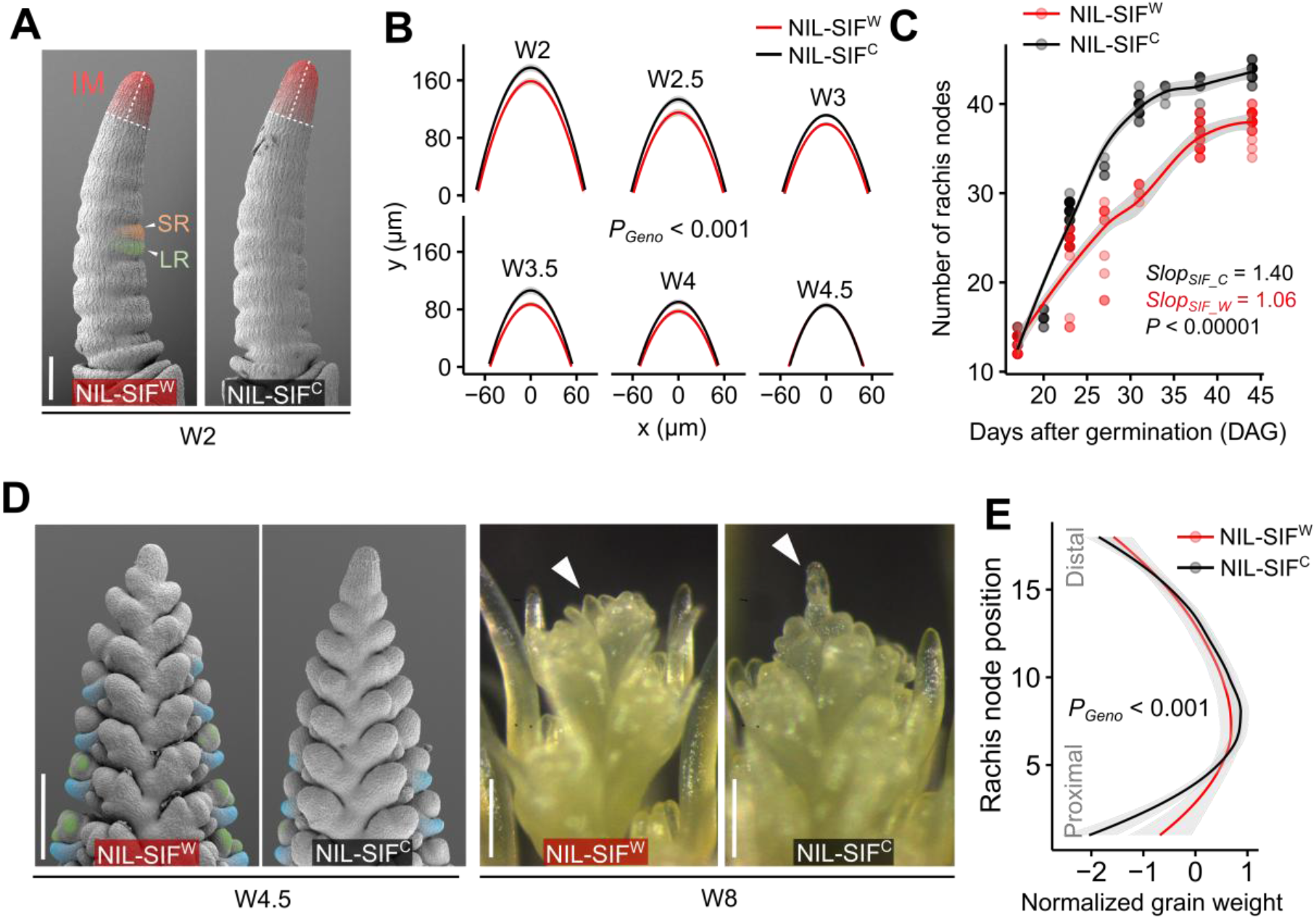
SIF-associated phenotypes in developing spikes. (A) Scanning electron microscopy (SEM) image showing inflorescence meristem (IM) at the W2 stage in NIL-SIF^C^ and NIL-SIF^W^ plants. IM, leaf-ridge (LR) and spikelet-ridge (SR) are marked with different colors. (B) IM morphology in NIL-SIF^C^ and NIL-SIF^W^ across stages. IM length and width were adjusted with a parabola function to improve visualization. (C) Comparison of the SM initiation rate between NIL-SIF^W^ and NIL-SIF^C^ plants. NIL-SIF^C^ plants initiate SMs faster, resulting in more SMs, as reflected by the steeper slope. (D) Spike morphology in NIL-SIF^C^ and NIL-SIF^W^ after the MYP stage. NIL-SIF^W^ plants exhibit better-developed distal floral organs at the W4.5 stage, and their IMs remain viable at W8. Lemma and stamen primordia are colored with cyan and green, respectively. (E) Grain yield distribution along the spikes of NIL-SIF^W^ and NIL-SIF^C^ at maturity. Scale bars: 200 µm in (A); 500 µm in (D).

**Extended Data Fig. 6.**
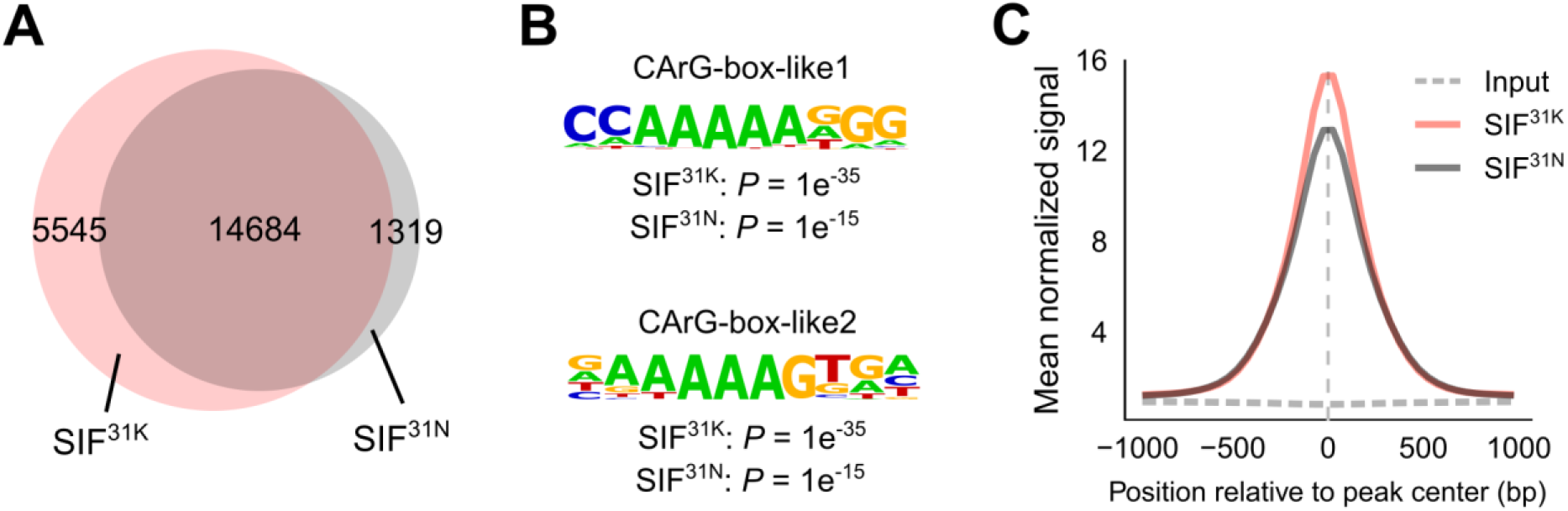
Functional significance of the K31N substitution on SIF determined by DAP-seq. (A) Venn diagram showing a greater number of binding peaks for SIF^31K^ than for SIF^31N^ revealed by DAP-seq. (B) Differential enrichment levels of two CArG-box-like motifs from the significant peaks bound by SIF^31K^ and SIF^31N^. (C) Normalized read coverage at the significant peaks bound by SIF^31K^ and SIF^31N^.

**Extended Data Fig. 7.**
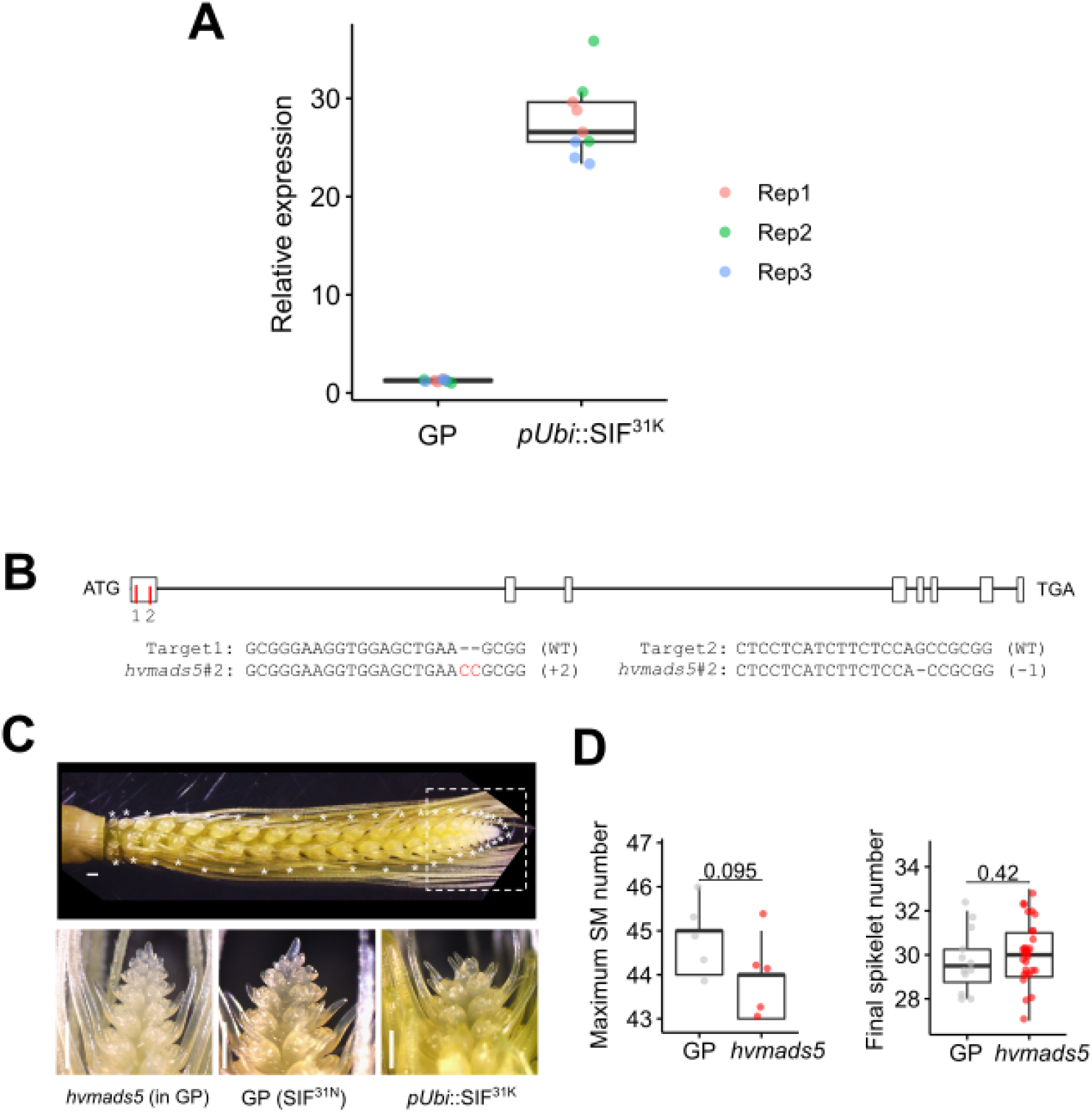
Functional validation of *SIF*. (A) Relative expression of *SIF* in the GP wild type and pUbi::SIF^31K^ transgenic line from W2 stage spikes, determined by RT-qPCR. Transcript abundance was normalized to Actin. Colored symbols indicate technical replicates grouped by biological replicate (n = 3 biological replicates, three technical replicates each). (B) *HvMADS5* (*SIF*) gene editing in Goden promise (GP). Schematic of the HvMADS5 gene structure and the corresponding *hvmads5*#2 mutant allele, redrawn from^36^. (C) Representative spike image at ∼W5.5 (MYP stage) illustrating how maximum SM number was quantified. Asterisks indicated individual rachis nodes, each subtending three SMs. The dashed box marks the growing spike apex, shown at higher magnification below for *hvmads5*, GP (SIF^31N^) and SIF^31K^ overexpression line (*pUbi*::SIF^31K^). <u>^36^</u>Scale bars: 500 µm. (D) Statistical comparisons of maximum SM number and final spikelet number in GP and its SIF/HvMADS5 edited plants. Not obvious morphological changes were found between the transgenic plants (*sif*/*hvmads5*) and the wild-type GP (SIF^31N^). Significant levels are determined by two-sided Student’s *t*-test.

**Extended Data Fig. 8.**
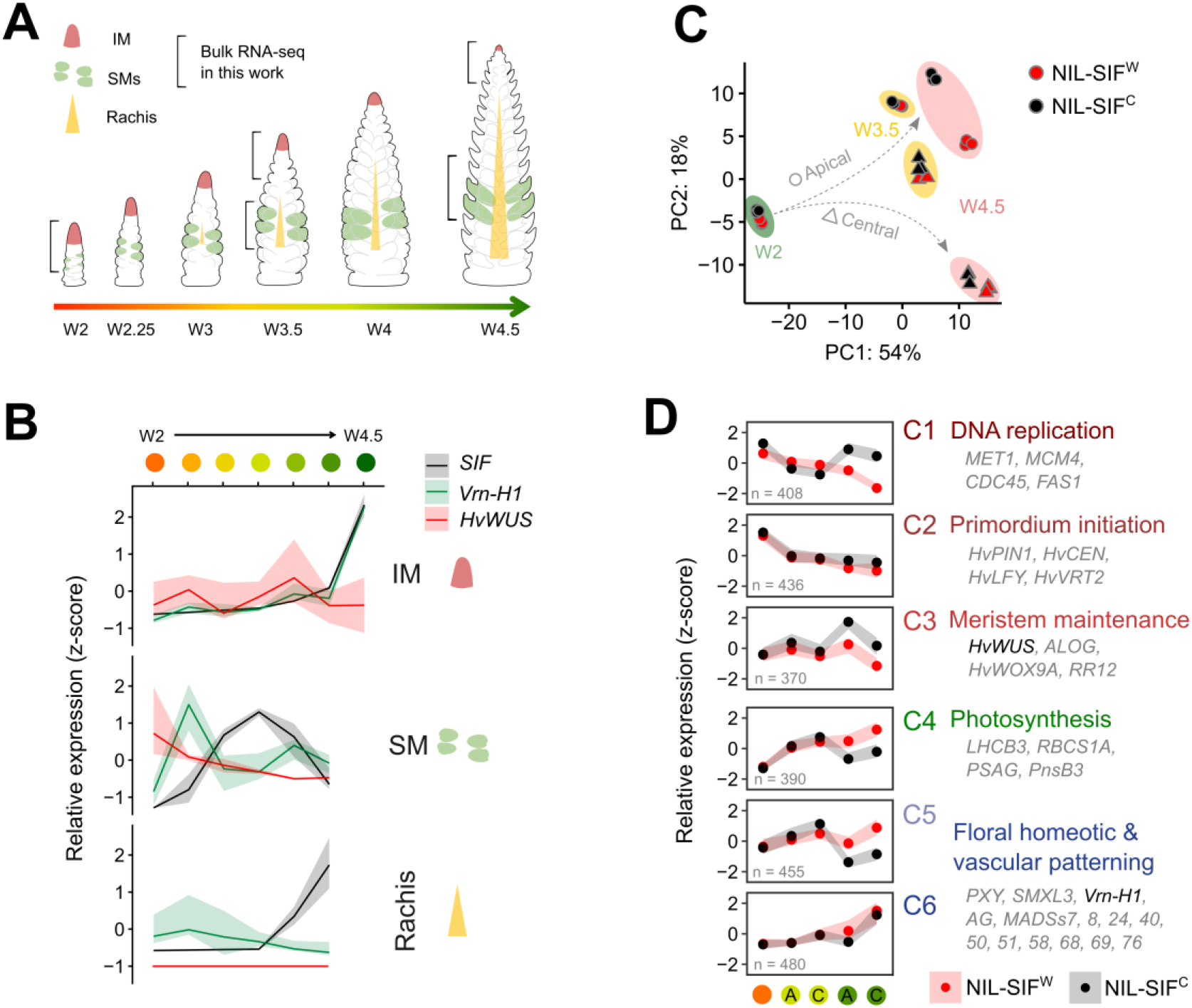
SIF reprograms the transcriptome during SM initiation and differentiation. (A) Schematic illustration of the spikelet organogenesis and the RNA-seq sampling strategy. The fates of the IM (red), SM (green), and central rachis (orange) are highlighted to illustrate their changing developmental fates throughout spike devleopment. Brackets indicate the three developmental stages and tissue sampled for bulk RNA-seq in this work. The colored arrow denotes the progression of spike development from W2 to W4.5, with the color gradient used consistently thourghout panel (D). (B) Gene expression (z-score) profiles for *HvWUS*, *Vrn-H1* and *SIF* during spikelet initiation and development across three tissue types. Source data from^47^. (C) Principal component analysis (PCA) of global gene expression profiles from the RNA-seq. Only the first two components were displayed. Samples were colored according to tissue type, developmental trajectory or genotype as in (A, color gradient of the arrow), which revealed a good coverage of the genotypic difference, developmental gradient or different tissue types. (D) Expression trajectories of DEGs relevant to spikelet organogenesis caused by *SIF* variation. Lines represent the mean scaled expression (row-wise z-score transformed TPMs) of genes within each cluster across spike developmental stages. Representative genes and enriched biological functions are shown for each cluster. Heterochronic shifts in the expression of genes controlling meristem maintenance, SM specification and vascularization were observed in NIL-SIF^C^ compared with NIL-SIF^W^. *HvWUS* and *Vrn-H1* were highlighted. See also Extended Data Table 4.

**Extended Data Fig. 9.**
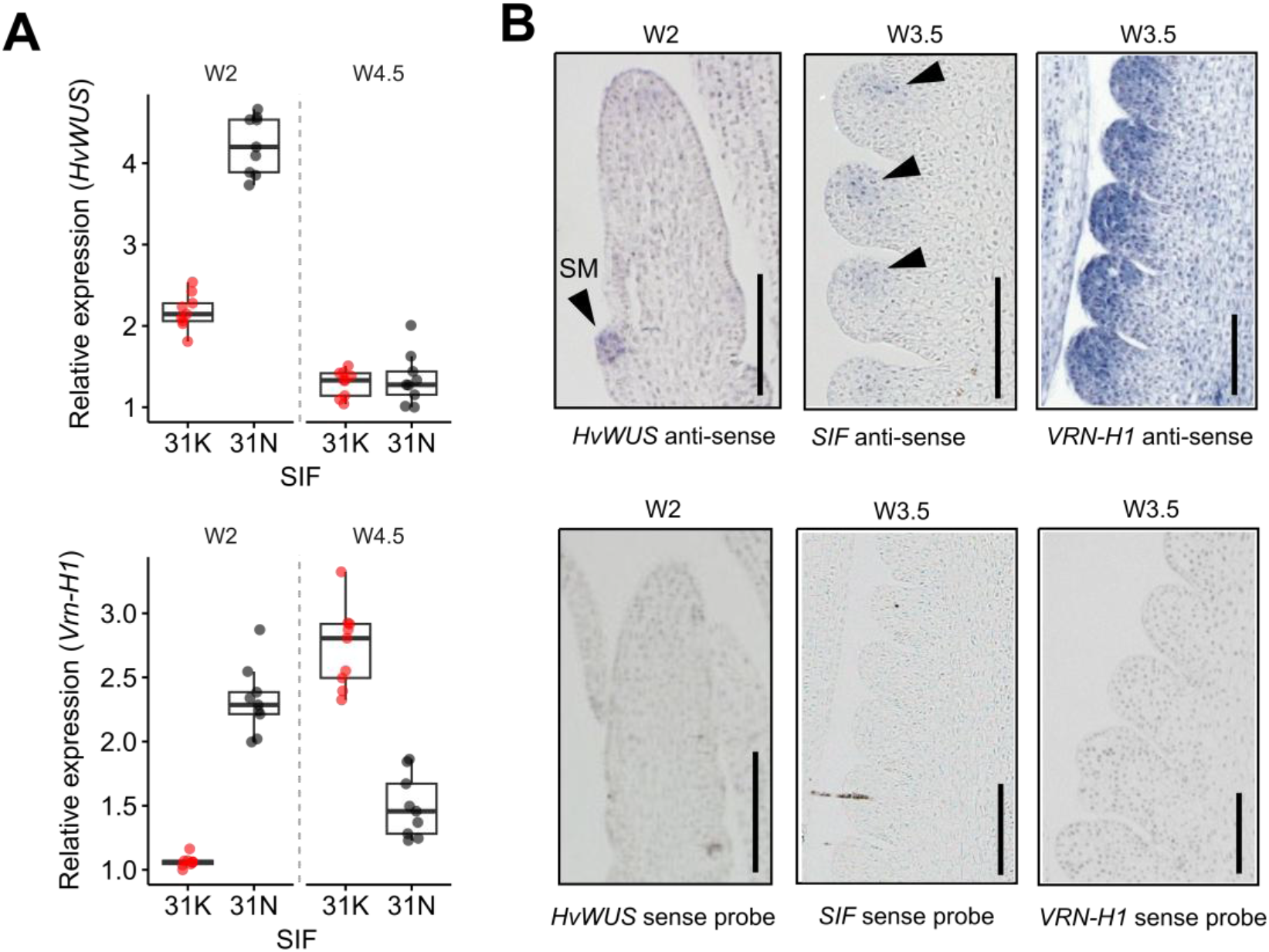
Expression of *HvWUS*, *SIF* and *Vrn-H1* during spike development. (A) Relative expression of *HvWUS* and *Vrn-H1* in developing spikes of NIL-SIF^C^ (31N) and NIL-SIF^W^ (31K) at W2 and W4.5 stages, determined by RT–qPCR. *HvWUS* expression is higher in the NIL-SIF^C^ background at W2, whereas *Vrn-H1* exhibits a heterochronic shift, with higher expression in the NIL-SIF^C^ background at W2 but higher expression in the NIL-SIF^W^ background at W4.5. (B) mRNA *in situ* hybridization of *HvWUS*, *SIF* and *Vrn-H1* in developing spikes. Barley developing spikes hybridized with sense probes from both genes were used as negative controls. Arrowheads indicate sites of transcript accumulation, with *Vrn-H1* expression exhibiting a broader expression domain throughout developing floral organs at W3.5. Scale bars: 200 µm.

**Extended Data Fig. 10.**
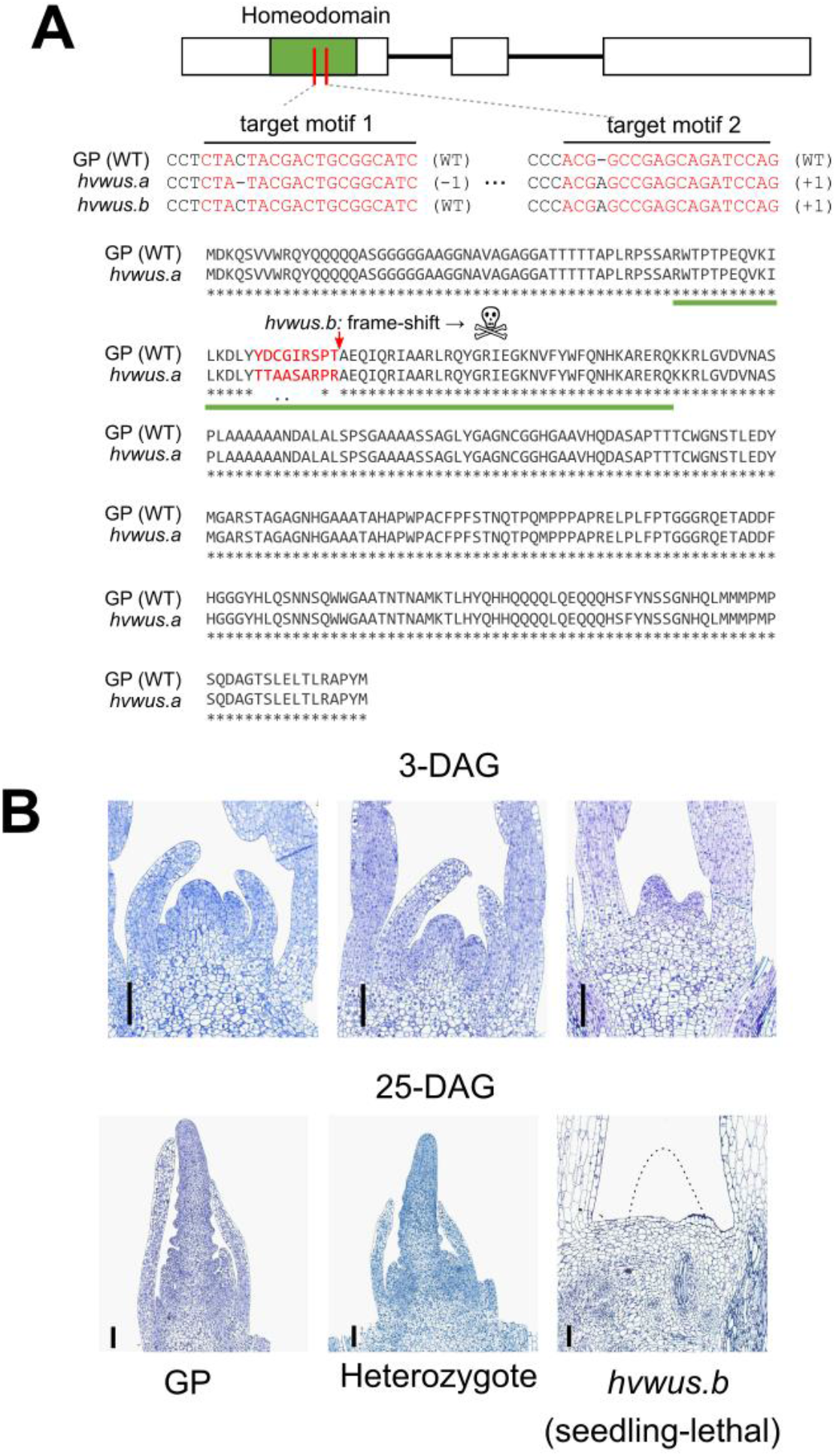
CRISPR-Cas9 knockout of *HvWUS*. (A) Schematic illustration of the two gRNAs and the mutations. Positions of the two target motifs (1 & 2) in the homeodomain (HB) for guide RNA design are indicated. The *hvwus.a* allele carries mutations in both gRNAs, causing an in-frame shifting of 9 amino acids within the HB domain, whereas the *hvwus.b* allele carries a mutation only at guide2, leading to a frameshift that disrupts the rest of the protein. (B) Resin sectioning of the developing apex in the *hvwus.b* mutant and Golden Promise (GP). No apical meristems were observed in the *hvwus.b* mutant at the seedling stage, leading to lethality at the 3-leaf stage. Heterozygote from a segregating T_3_ generation was included as a control. DAG, days after germination. Scale bars: 100 µm.

**Extended Data Fig. 11.**
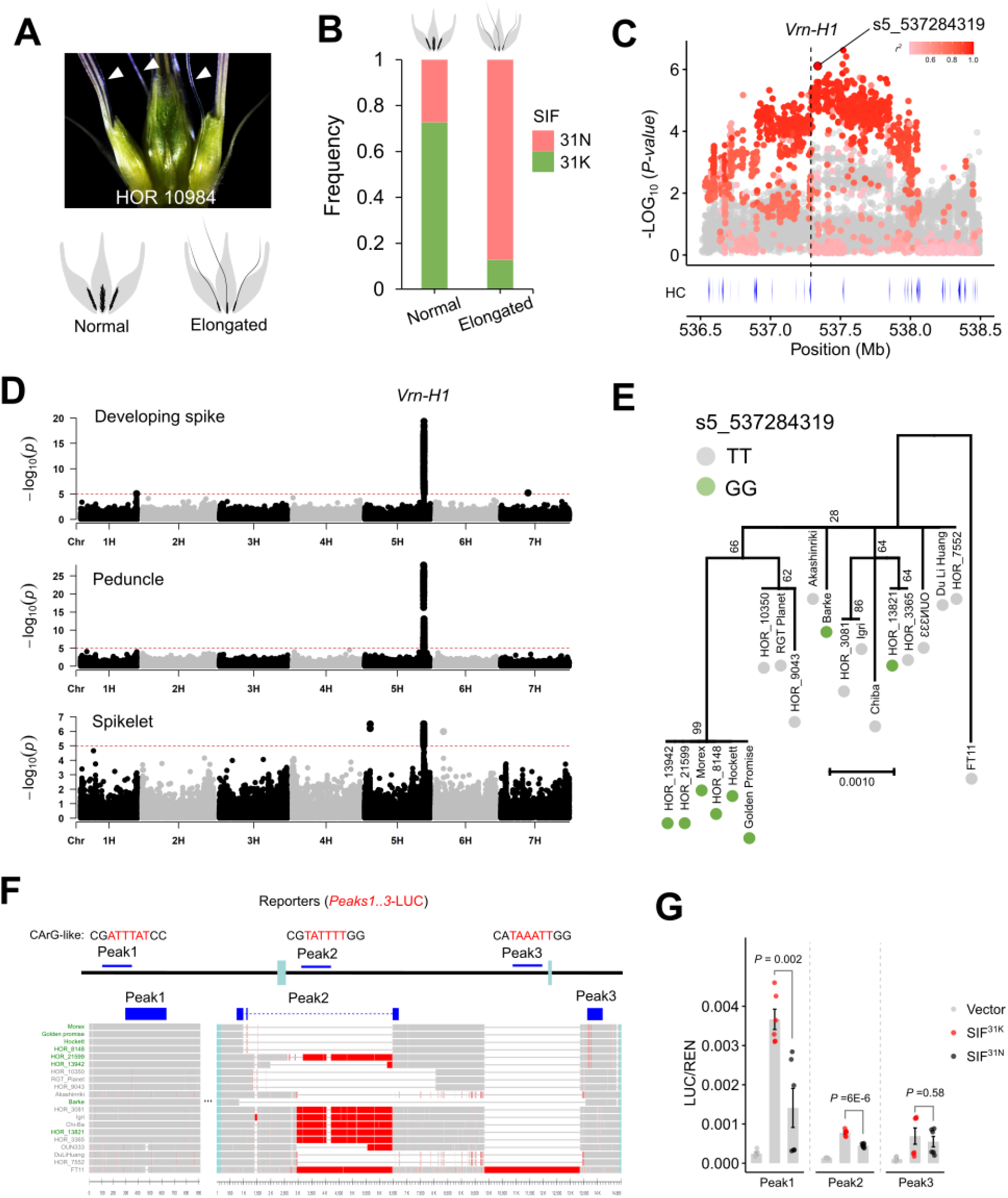
Genetics of rachilla elongation associated with *Vrn-H1* intron variation and expression. (A) A representative image showing the rachilla elongation phenotype in HOR_10984. Schematic illustration of spikelets with normal versus elongated rachilla is shown below. (B) SIF K31N variation is linked to the variation in rachilla length in the D6S population. Note that barley accessions carrying the SIF^31K^ allele produced shorter rachillas compared to those with SIF^31N^ allele. (C) Local Manhattan of the *Vrn-H1* locus for rachilla elongation. Surrounding SNPs were colored according to their linkage relationship (*r^2^*) with the leading SNP (s5_537518593). Note that SNP s5_537284319 (Morex V2) corresponds to the leading SNP (S5H_532318701, Barke V1) identified in expression associations for *Vrn-H1*. (D) Expression association analysis for *Vrn-H1*. *Vrn-H1* TPMs from three tissues were used as quantitative phenotypes for the association mapping. A major *cis*-eQTL was identified at the Vrn-H1 locus in developing spikes and peduncles, but was weaker in spikelets. The association peak (leading SNP: S5H_532318701) also accounts for the variation in rachilla elongation in the D6S population. Red dashed horizontal lines represent the Bonferroni-corrected significance threshold. Source data from^97^. (E) Neighbor joining phylogeny of the *Vrn-H1* genomic region in the 20 barley reference genomes^34^. Barley accessions were colored according to the leading SNP s5_537518593. Bootstrap values for each node are included (in percentage). (F) Association peaks for rachilla elongation and *Vrn-H1* expression are linked with natural variations within the SIF binding peaks (Peak1-Peak3). Alignments highlighted in red indicate sequence variation among the 20 reference genomes. Note that large insertions/deletions were observed within the second SIF binding peak within the first intron, whereas the first and third peak primarily harbored SNPs or short deletions. CArG-like motifs identified near the binding peaks were highlighted. (G) Transient Dual-LUC assays testing the regulatory activity of SIF on three DAP-seq binding regions within the *Vrn-H1* locus. Three regulatory fragments (Peak1–Peak3) containing SIF-binding sites were fused to the LUC reporter and co-expressed with either SIF^31K^ or SIF^31N^. Peak1 and Peak2 showed significantly stronger reporter activation by SIF^31K^ than by SIF^31N^, whereas no significant difference was detected for Peak3. Peak1–Peak3 co-incubated with an empty vector were served as internal controls for their respective groups. Error bars represent s.e.m.; individual biological replicates are shown as dots. *P* values were determined by two-sided Student’s *t*-tests.

**Extended Data Fig. 12.**
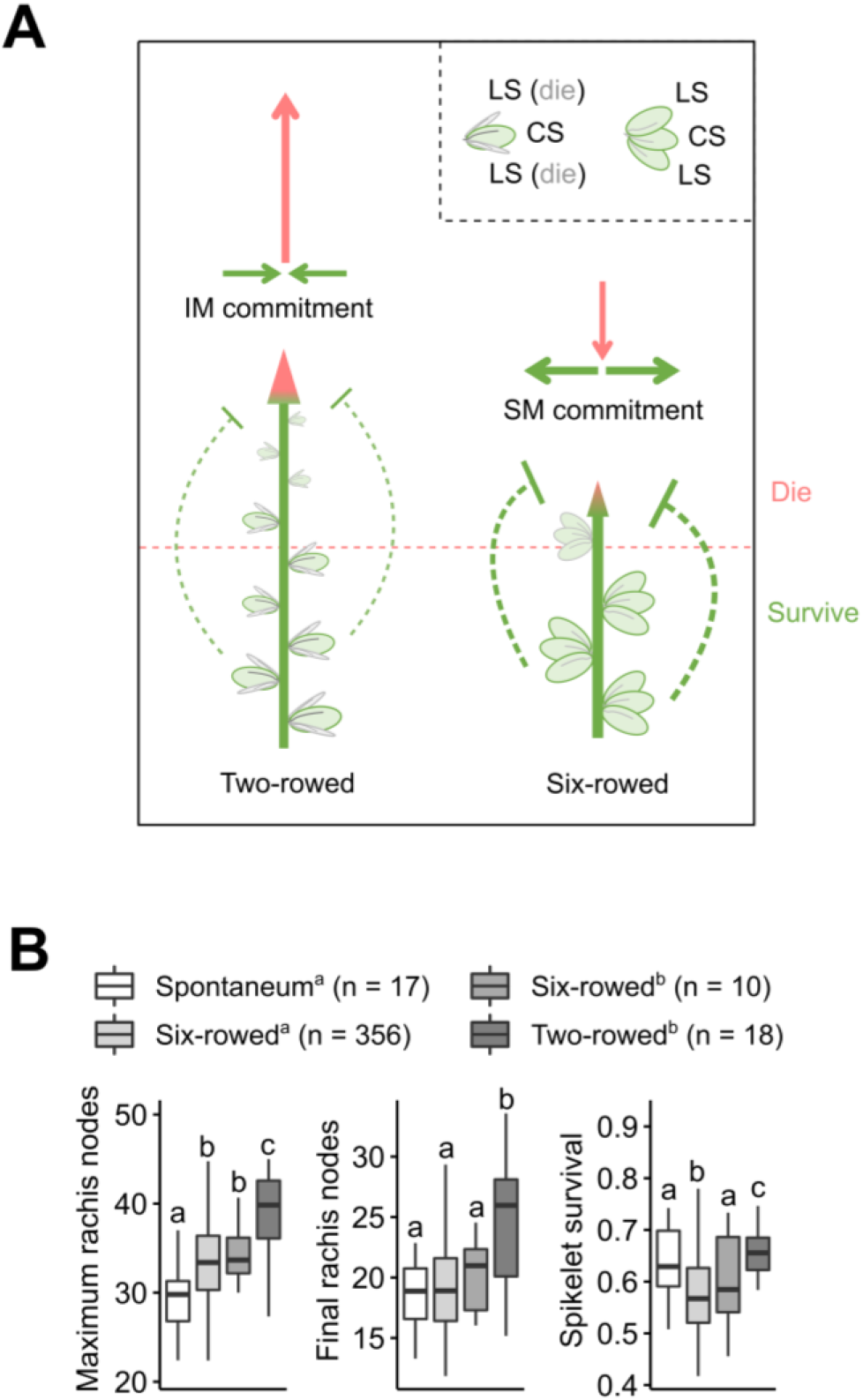
Diversified reproductive strategies between row-types. (A) Schematic illustration of the reproductive strategies favored by different row-types. Two-rowed spikes abort two lateral florets at early developmental stages, reducing demands for SM commitment and enhancing IM commitment, which leads to increased SM initiations but more distal spikelet degeneration. By contrast, six-rowed spikes place higher demands on SM commitment (e.g., three times higher), potentially resulting in reduced SM initiation while maintaining higher survival. (B) Comparison of maximum SM number (recoded as rachis nodes), final rachis node and spikelet survival among different barley subpopulations. Six-rowed barleys do not yield three times than two-rowed barleys due to reduced IM indeterminacy (lower SM initiation). Letters above boxplot indicate statistical significance corresponding to Tukey’s HSD (α = 0.05). Letters from the legend indicates source of the phenotypic data. a:^30^; b:^60^.

**Extended Data Fig. 13.**
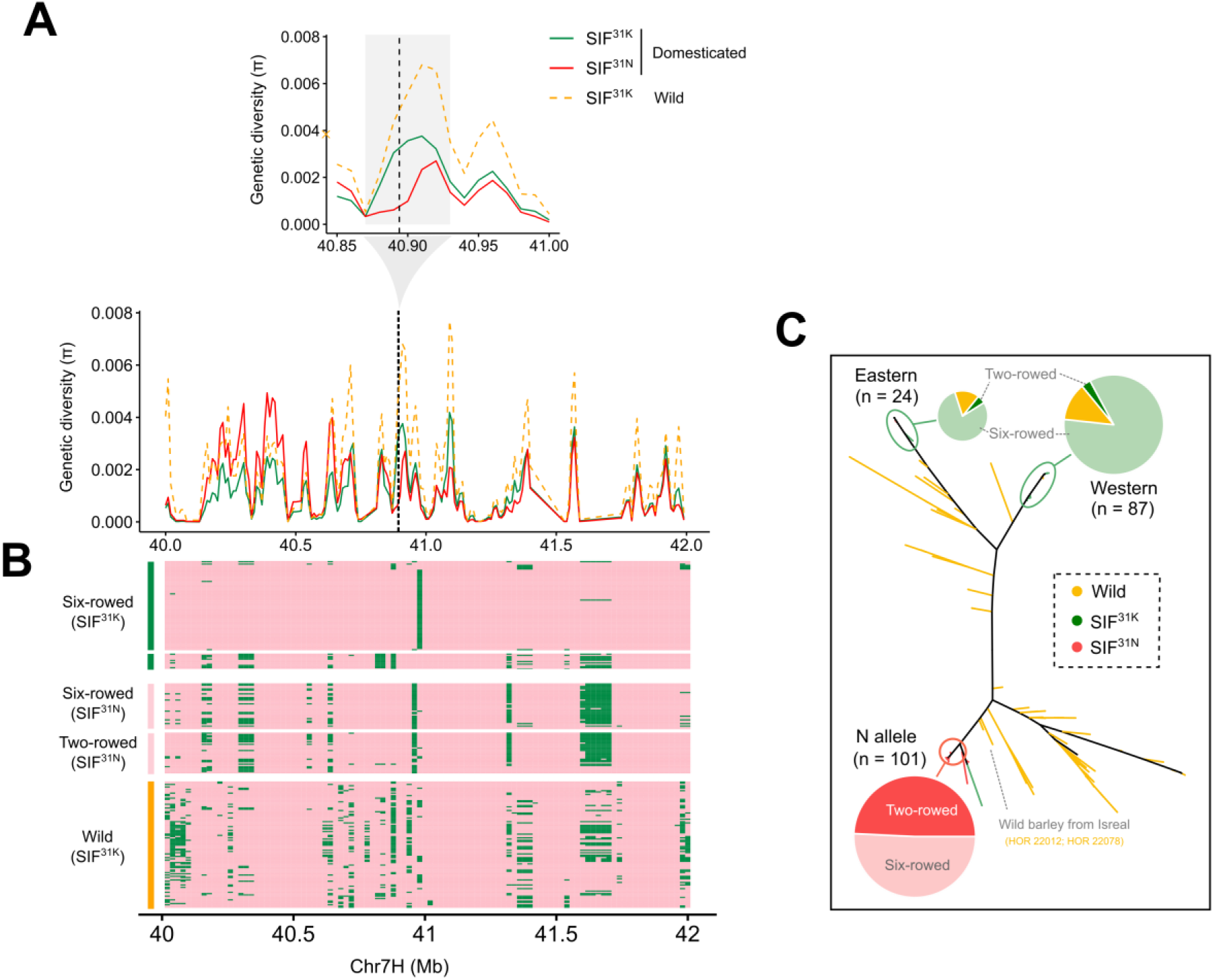
*SIF* selection footprint. (A) Genetic diversity at the *SIF* locus. A genomic region (∼60 Kb) with relatively low diversity in SIF^31N^ is highlighted in grey. The position of the *SIF* gene is indicated with a black dashed line. Source data from from^34^. (B) Genomic structure surrounding the *SIF* locus. Major and minor alleles on each polymorphic site are represented in green and pink, respectively (see methods). The genotypes with different *SIF* alleles, row-types or domestication status are shown on the left. Source data from from^34^. (C) Neighbor-joining clustering of the *SIF* locus based on 10,220 SNPs within the 1-Mb genomic region surrounding *SIF*. Allelic frequencies among subpopulations remained consistent with the analysis of an independent dataset shown in Fig. 5A. The analysis further revealed at least two independent origins of the SIF^31K^ allele in domesticated barley, whereas the SIF^31N^ allele likely originated from a single domestication event during low-latitudinal (or high temperature) adaptation. The most closely related wild counterparts for SIF^31N^ haplotype from Israel are highlighted. Source data from^34^.

**Extended Data Fig. 14.**
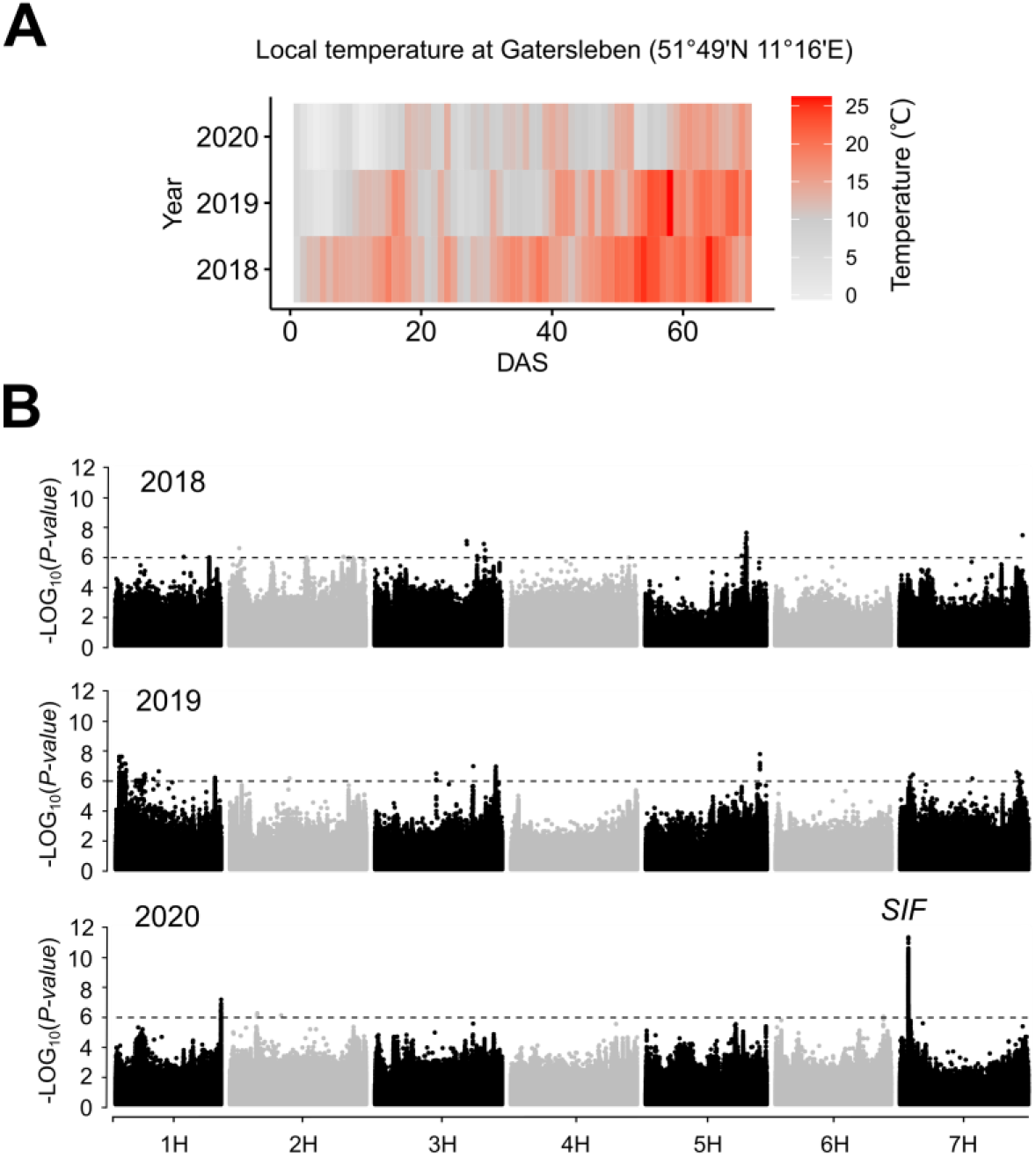
Environmental sensitivity of *SIF* on SM initiation rate revealed by GWAS in a three-year common-garden experiment. (A) Local temperature between 2018-2020 at the Gatersleben (related to Fig. 5D). Local temperature was recorded daily from a local weather station. The critical period for spikelet initiation and development ranges from 20 to 50 days after sowing (DAS). (B) GWAS of SM initiation rate across three constitutive years (2018-2020) of field trial in the D6S population (related to Fig. 5D). Dashed horizontal lines represent the Bonferroni-corrected significance threshold.

**Extended Data Fig. 15.**
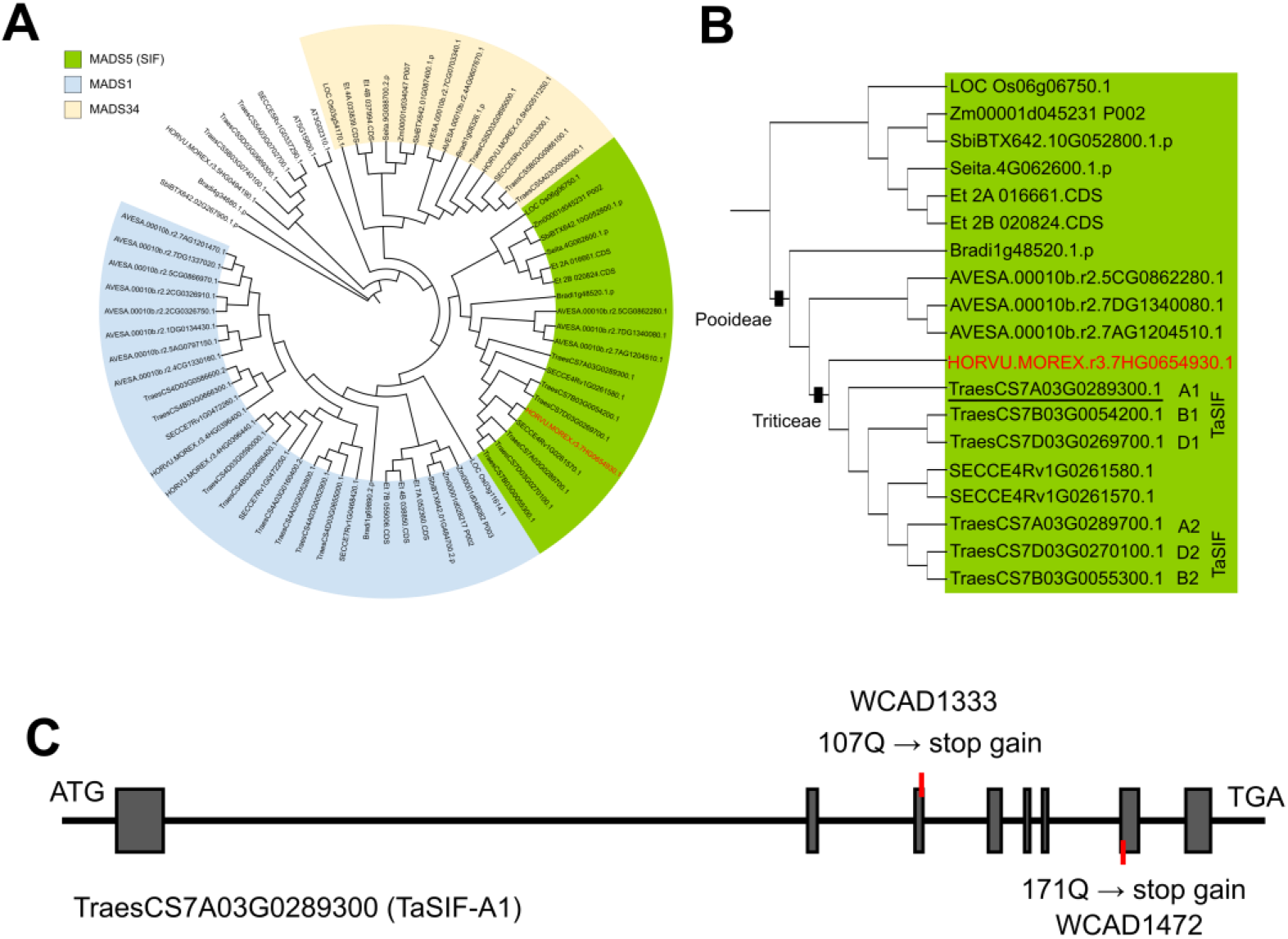
Phylogenetic analysis of the LOFSEP-clade MADS-box genes in grasses. (A) A maximum-likelihood (ML) phylogenetic tree of the LOFSEP-clade MADS-box proteins among 10 plant species. 71 protein sequences from rye (*Secale cereale*), wheat (*Triticum aestivum*), barley (*Hordeum vulgare*), Brachypodium (*Brachypodium distachyon*), oat (*Avena sativa*), sorghum (*Sorghum bicolor*), maize (*Zea mays*), rice (*Oryza sativa*), millet (*Setaria italica*) and Arabidopsis (*Arabidopsis thaliana*) were used to build the tree. The grass-specific LOFSEP clade is highlighted with different colors, with barley SIF (MADS5) marked in red. Note that tandem gene duplications in the MADS1 and MADS5 clades were identified in the Triticeae tribe. Protein sequences were downloaded from Phytozome (https://phytozome-next.jgi.doe.gov/). (B) Close-up view of the MADS5 (SIF) clade. The closest Wheat SIF ortholog (TraesCS7A03G0289300, designed as TaSIF-A1) was underlined. Barley SIF was indicated in red. (C) Schematic illustration of two mutant alleles carrying stop-gain mutations in *TaSIF-A1* within the Cadenza background (related to Fig. 6E).

